# Associations between locus coeruleus MRI contrast and physiological responses to acute stress in younger and older adults

**DOI:** 10.1101/2022.03.12.484104

**Authors:** Shelby L. Bachman, Kaoru Nashiro, Hyunjoo Yoo, Diana Wang, Julian F. Thayer, Mara Mather

## Abstract

Acute stress activates the brain’s locus coeruleus (LC)-noradrenaline system. Recent studies indicate that a magnetic resonance imaging (MRI)-based measure of LC structure is associated with better cognitive outcomes in later life. Yet despite the LC’s documented role in promoting physiological arousal during acute stress, no studies have examined whether MRI-assessed LC structure is related to arousal responses to acute stress. In this study, 102 younger and 51 older adults completed an acute stress induction task while we assessed multiple measures of physiological arousal (heart rate, breathing rate, systolic and diastolic blood pressure, sympathetic tone, and heart rate variability, HRV). We used turbo spin echo MRI scans to quantify LC MRI contrast as a measure of LC structure. We applied univariate and multivariate approaches to assess how LC MRI contrast was associated with arousal at rest and during acute stress reactivity and recovery. In older participants, having higher caudal LC MRI contrast was associated with greater stress-related increases in systolic blood pressure and decreases in HRV, as well as lower HRV during recovery from acute stress. These results suggest that having higher caudal LC MRI contrast in older adulthood is associated with more pronounced physiological responses to acute stress. Further work is needed to confirm these patterns in larger samples of older adults.

## 1. Introduction

Acute stress is a recurring feature of daily life, occurring in response to various psychosocial and environmental stressors. The nervous system’s highly-conserved response to acute stress is designed to promote behaviors and processes that facilitate survival in the face of such stressors (Johnson et al., 1992; Monaghan & Spencer, 2014). At the same time, there is substantial variability in how individuals respond to stress (Rab & Admon, 2021; Sapolsky, 2015; Zänkert et al., 2019), with exaggerated or prolonged stress responses associated with adverse health outcomes. Individuals with excessive responses to and impaired recovery from acute stress are at elevated risk for atherosclerosis, hypertension, myocardial infarction, and cardiovascular disease mortality (Chida & Steptoe, 2010; Panaite et al., 2015; Treiber et al., 2003). Furthermore, stress contributes to dementia risk (Justice, 2018; Lyons & Bartolomucci, 2020; Yuede et al., 2018), which may in part be mediated by noradrenergic modulation of β-amyloid and tau production and clearance (Mather, 2021). Thus, characterizing factors that may protect against stress vulnerability across the adult lifespan is an important aim of psychophysiological research.

Acute stress responses engage both the hypothalamic-pituitary-axis and the brain’s noradrenergic system, the hub of which is the locus coeruleus. The locus coeruleus (LC) is a nucleus within the pons of the brainstem that releases norepinephrine throughout the brain and spinal cord (Dahlström & Fuxe, 1964; Schwarz & Luo, 2015). Noradrenergic projections from the LC reach cortical regions implicated in attention, learning and memory (Sara, 2009), but the LC also sends projections to preganglionic sympathetic neurons in the spinal cord, which coordinate peripheral arousal responses (Jones & Yang, 1985). Besides releasing norepinephrine to the brain and spinal cord, the LC is innervated by brain regions including the central nucleus of the amygdala, the paraventricular nucleus of the hypothalamus, and the nucleus paragigantocellularis (Aston-Jones et al., 1986; Curtis et al., 2002; Mather, 2020; Samuels & Szabadi, 2008; Van Bockstaele & Aston-Jones, 1995). These inputs provide visceral feedback signals which the LC integrates to adaptively regulate norepinephrine release and, in turn, arousal levels (Morris et al., 2020a).

As an arousal hub region within the nervous system, the LC is robustly activated in response to diverse stressors (Morilak, 2007; Valentino & Van Bockstaele, 2008). During acute stress, in tandem with hypothalamic-pituitary-adrenal (HPA) axis activation, corticotropin-releasing factor is released on the LC by the paraventricular nucleus, the central nucleus of the amygdala, and the bed nucleus of the stria terminalis (Johnson et al., 1992; Valentino & Van Bockstaele, 2005). Corticotropin-releasing factor increases the rate of tonic, or basal, norepinephrine discharge by LC neurons while decreasing the frequency of phasic, stimulus-evoked responses (Curtis et al., 1997; Valentino & Foote, 1988). A shift to higher tonic LC activity increases cortical levels of norepinephrine (Kawahara et al., 2000) and promotes adaptive behavioral and physiological shifts that subserve threat detection and avoidance, such as the reorienting of attention and cardiovascular reactivity (Bremner et al., 1996; Sara & Bouret, 2012; Wood & Valentino, 2017).

The LC’s short-term response to acute stress is adaptive, promoting behaviors that complete the stress cycle. Yet stress experienced over the longer term may have maladaptive consequences for the LC. Corticotropin-releasing factor exposure due to chronic stress causes morphological changes to LC neurons, increasing both dendritic arborization and the number of primary processes (Borodovitsyna et al., 2018). Stress also affects the activity of LC neurons, with LC neurons from rodents exposed to chronic stress exhibiting higher excitability and sensitivity relative to those from controls, as well as anxiety-like behaviors (Jedema & Grace, 2003; Mana & Grace, 1997; McCall et al., 2015). Together, these findings suggest that structure of the LC may be closely intertwined with autonomic responses to acute stress.

Despite the LC’s involvement in the stress response, little is known about how LC structure is related to physiological responses in humans. This may be due in part to limitations of studying the LC in vivo in humans due to the LC’s small size and location. Recently, the development of specialized MRI protocols, including high-resolution turbo spin echo (TSE) and magnetization transfer sequences (Betts et al., 2019b; Sasaki et al., 2006), has made quantifying LC structure in vivo possible. In these sequences, the LC appears as hyperintense regions bordering the fourth ventricle; signal intensity of the LC relative to that of surrounding pontine tissue, which will henceforth be referred to as LC MRI contrast, is thought to reflect LC structure (Keren et al., 2009). A recent study using such a protocol found that LC volume was higher in younger adults with anxiety disorders relative to healthy controls, and that, across the sample of younger adults, LC volume was positively correlated with levels of anxious arousal and general distress (Morris et al., 2020b). In separate studies, younger and older adults with higher LC MRI contrast had lower heart rate variability (HRV), a measure of parasympathetic control over heart rate, during a fear conditioning task (Mather et al., 2017), and younger adults with higher MRI contrast of the LC’s caudal region had lower average cortical thickness (Bachman et al., 2021).

Both HRV and cortical thickness are lower in individuals with stress- and anxiety-related disorders relative to healthy controls (Chalmers et al., 2014; Molent et al., 2018), and dysregulation of noradrenergic signaling is feature of such disorders (Hendrickson & Raskind, 2016; Ressler & Nemeroff, 2000). Together, this evidence suggests that in younger adults, LC structure may be associated with poorer stress- and anxiety-related outcomes. Yet there has been little work directly examining associations between LC structure and comprehensive physiological responses to acute stress, despite the LC’s projections to and innervation from sympathetic and parasympathetic arousal centers.

In contrast to the reports described above, studies of older adults have indicated that having higher LC MRI contrast is associated with better cognitive performance across domains (Dahl et al., 2019; Liu et al., 2020), higher cortical thickness (Bachman et al., 2021), and lower risk of developing mild cognitive impairment (Elman et al., 2021a) in older adulthood. The LC is the first brain region where tau pathology accumulates in the progression of Alzheimer’s disease (Braak et al., 2011), and older adults with Alzheimer’s disease have lower LC MRI contrast relative to healthy controls (Betts et al., 2019a; Takahashi et al., 2015). On the surface, these seemingly discrepant findings – LC structure being associated with better outcomes later in adulthood but poorer outcomes earlier in adulthood – suggest that LC structure may be more influenced by stress in younger adults and more by neurodegeneration in older adults. However, just as for younger adults, no studies to date have assessed whether LC structure in older adulthood is related to aspects of the parasympathetic and sympathetic response to stress.

In the present study, we attempted to fill these gaps in the literature by examining how responses to acute stress were related to LC MRI contrast in a sample of 102 younger and 51 older adults. Participants completed an acute stress induction protocol, and we assessed multiple measures of physiological arousal during rest, acute stress, and acute stress recovery. TSE MRI scans were collected to assess LC contrast along the LC’s rostrocaudal extent. Pairwise correlation and partial least squares correlation analyses were applied to assess how LC contrast was associated with multiple measures of physiological arousal in each age group. In line with previous findings of LC volume being positively correlated with levels of anxious arousal in younger adults and LC contrast being associated with lower parasympathetic control over heart rate (Mather et al., 2017; Morris et al., 2020b), we predicted that younger adults with higher LC contrast would have higher-magnitude responses to acute stress. In terms of predictions for older adults, in line with studies linking MRI-assessed LC structure to better cognitive and neural outcomes in aging (Dahl et al., 2019; Elman et al., 2021a; Liu et al., 2020), we originally expected that older adults with higher LC contrast would have lower-magnitude acute stress responses, reflecting reduced potential impacts of stress. However, another perspective is that acute stress reactivity in aging is beneficial, reflecting a responsive, flexible autonomic system in the context of normal sympathetic tone. From this perspective, a competing possibility was that older adults with higher LC contrast would have larger-magnitude acute stress responses.

## 2. Results

### 2.1. Effectiveness of the acute stress induction protocol

Average measures of arousal during the baseline, challenge and recovery phases of the stress induction task are presented in Figure 1. Linear mixed-effects analyses were used to assess whether during the acute stress induction protocol, average measures of arousal differed during the baseline and challenge phases, and during the challenge and recovery phases (Table 1). Results of all planned, pairwise comparisons of each measure for each phase contrast and age group are presented in the Supplementary Results (Section 1). For heart rate, breathing rate, systolic blood pressure, and diastolic blood pressure, we found significant elevations from the baseline to the challenge phase (*p*s <= 0.012; Table 1), and significant decreases from the challenge to the recovery phase (*p*s < .001; Table 1). Sympathetic tone did not increase significantly from baseline to the challenge phase (*p* = .244; Table 1) but decreased significantly from challenge to recovery (*p* = .002; Table 1). For systolic blood pressure, we found a significant phase (recovery-challenge) x age group interaction (*p* = .006), which was driven by greater challenge-to-recovery decreases in blood pressure for older compared to younger participants, although challenge-to-recovery changes were significant in both age groups (Supplementary Results, Section 1).

**Figure 1.**
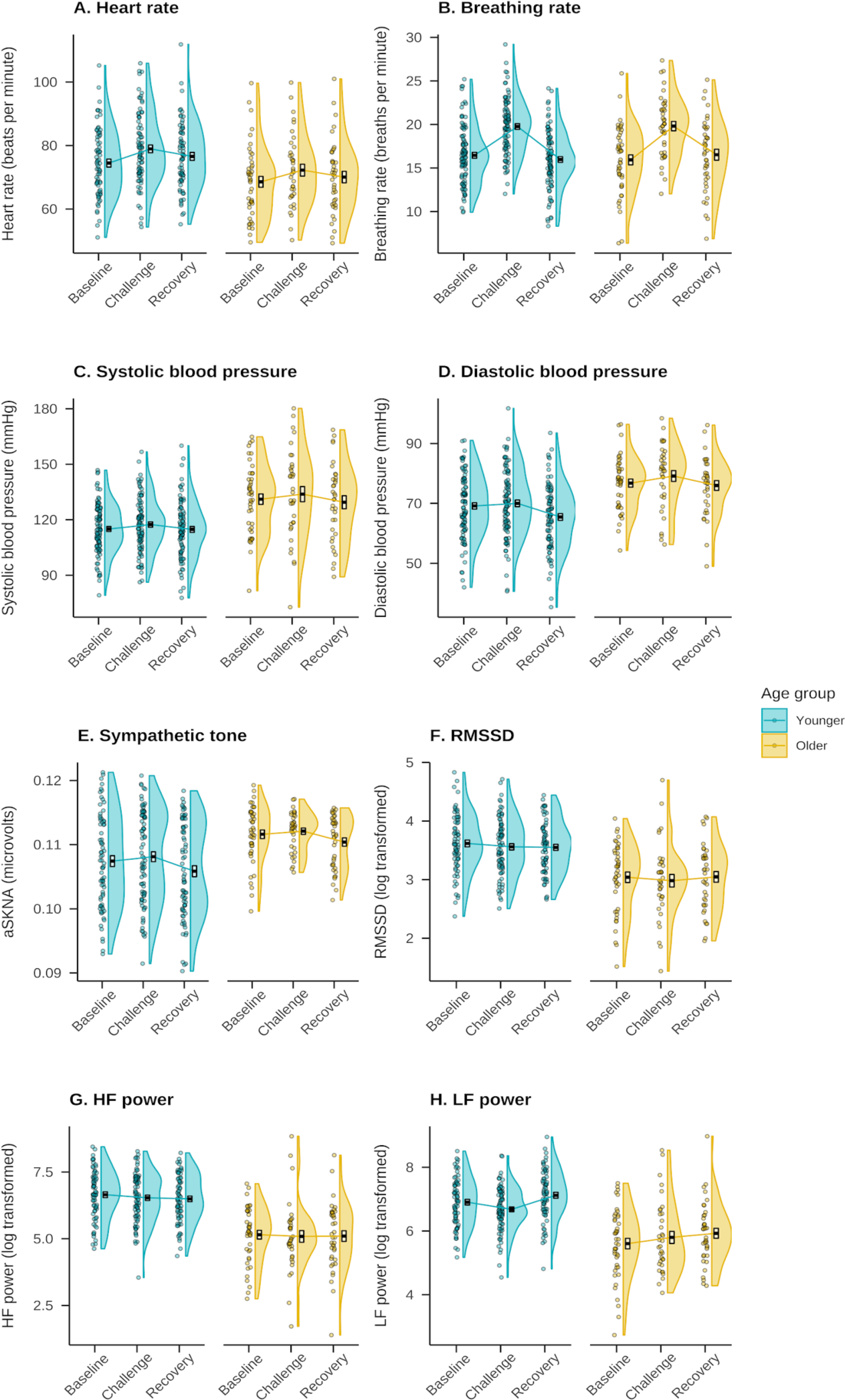
Average measures of physiological arousal during each phase of the stress induction protocol. Crossbars reflect standard errors of the mean. LF = low-frequency; HF = high-frequency; RMSSD = root mean square of the successive differences.

**Table 1.**
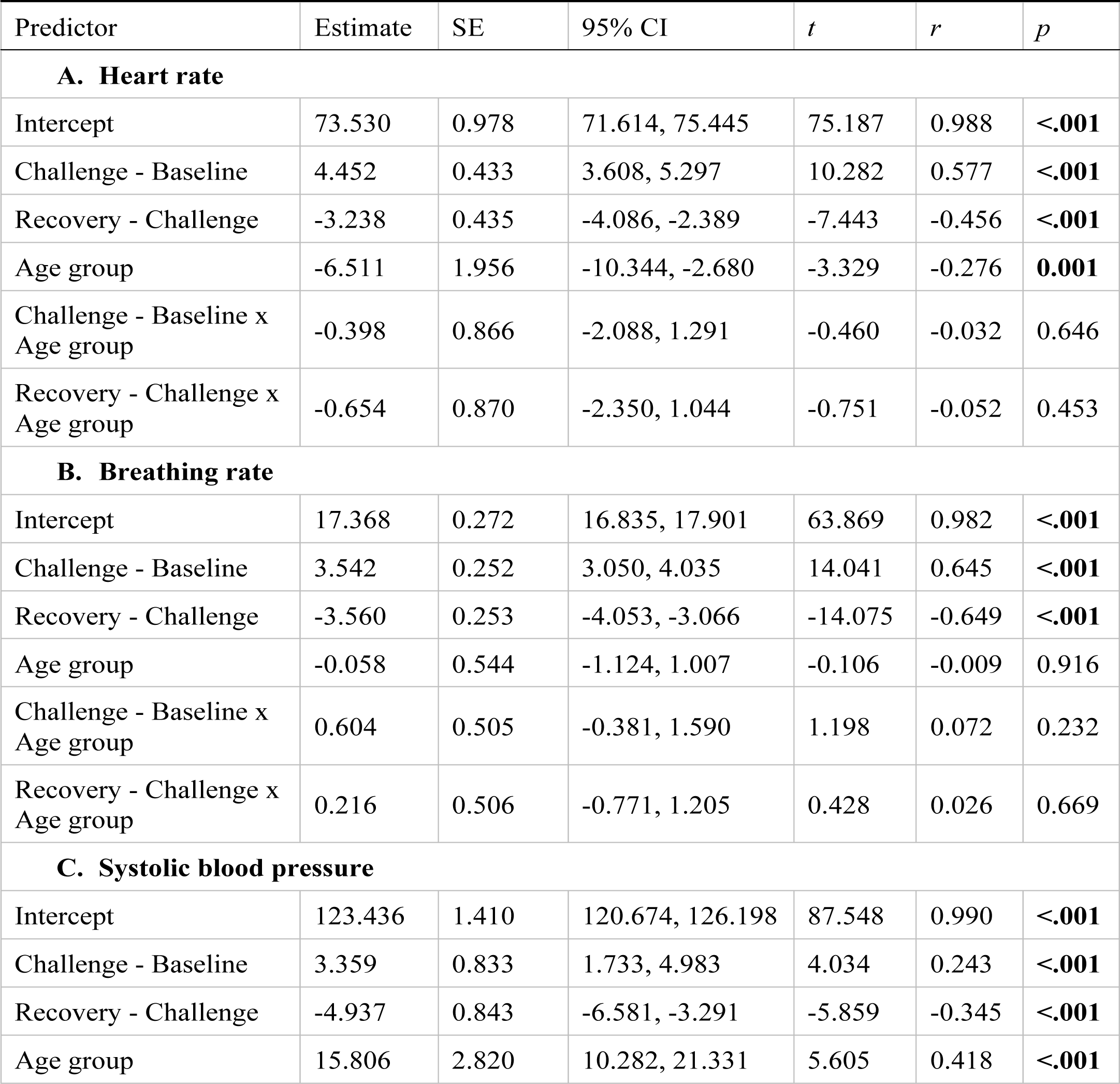
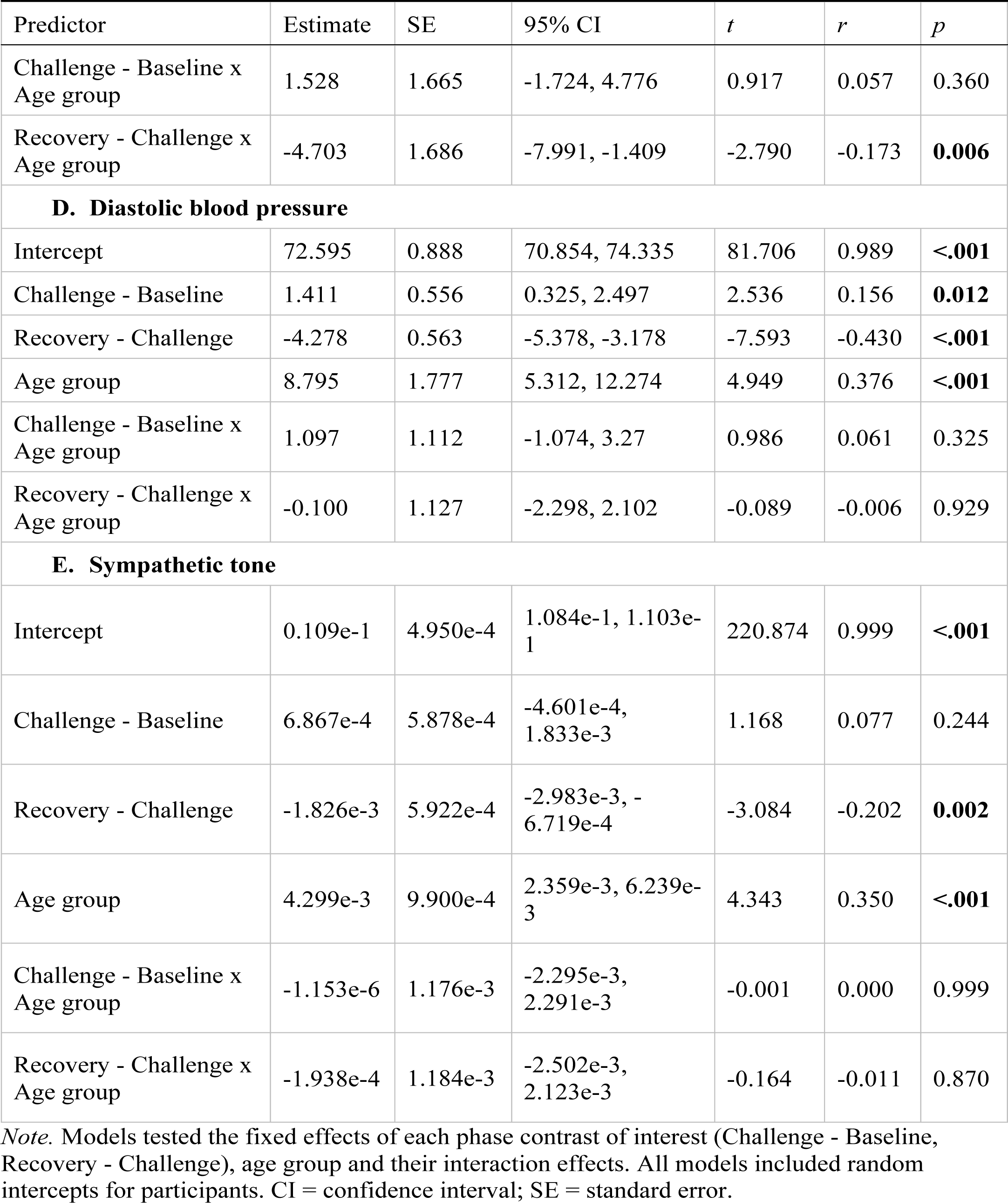
Results of linear mixed effects analyses testing that during the acute stress induction protocol, average heart rate (A), breathing rate (B), systolic blood pressure (C), diastolic blood pressure (D), and sympathetic tone (E) differed during the baseline and challenge phases and during the challenge and recovery phases.

| Predictor | Estimate | SE | 95% CI | *t* | *r* | *p* |
| --- | --- | --- | --- | --- | --- | --- |
| **A. Heart rate** |  |  |  |  |  |  |
| Intercept | 73.530 | 0.978 | 71.614, 75.445 | 75.187 | 0.988 | <.001 |
| Challenge - Baseline | 4.452 | 0.433 | 3.608, 5.297 | 10.282 | 0.577 | <.001 |
| Recovery - Challenge | -3.238 | 0.435 | -4.086, -2.389 | -7.443 | -0.456 | <.001 |
| Age group | -6.511 | 1.956 | -10.344, -2.680 | -3.329 | -0.276 | 0.001 |
| Challenge - Baseline x Age group | -0.398 | 0.866 | -2.088, 1.291 | -0.460 | -0.032 | 0.646 |
| Recovery - Challenge x Age group | -0.654 | 0.870 | -2.350, 1.044 | -0.751 | -0.052 | 0.453 |
| **B. Breathing rate** |  |  |  |  |  |  |
| Intercept | 17.368 | 0.272 | 16.835, 17.901 | 63.869 | 0.982 | <.001 |
| Challenge - Baseline | 3.542 | 0.252 | 3.050, 4.035 | 14.041 | 0.645 | <.001 |
| Recovery - Challenge | -3.560 | 0.253 | -4.053, -3.066 | -14.075 | -0.649 | <.001 |
| Age group | -0.058 | 0.544 | -1.124, 1.007 | -0.106 | -0.009 | 0.916 |
| Challenge - Baseline x Age group | 0.604 | 0.505 | -0.381, 1.590 | 1.198 | 0.072 | 0.232 |
| Recovery - Challenge x Age group | 0.216 | 0.506 | -0.771, 1.205 | 0.428 | 0.026 | 0.669 |
| **C. Systolic blood pressure** |  |  |  |  |  |  |
| Intercept | 123.436 | 1.410 | 120.674, 126.198 | 87.548 | 0.990 | <.001 |
| Challenge - Baseline | 3.359 | 0.833 | 1.733, 4.983 | 4.034 | 0.243 | <.001 |
| Recovery - Challenge | -4.937 | 0.843 | -6.581, -3.291 | -5.859 | -0.345 | <.001 |
| Age group | 15.806 | 2.820 | 10.282, 21.331 | 5.605 | 0.418 | <.001 |
| Challenge - Baseline x Age group | 1.528 | 1.665 | -1.724, 4.776 | 0.917 | 0.057 | 0.360 |
| Recovery - Challenge x Age group | -4.703 | 1.686 | -7.991, -1.409 | -2.790 | -0.173 | **0.006** |
| **D. Diastolic blood pressure** |  |  |  |  |  |  |
| Intercept | 72.595 | 0.888 | 70.854, 74.335 | 81.706 | 0.989 | **<.001** |
| Challenge - Baseline | 1.411 | 0.556 | 0.325, 2.497 | 2.536 | 0.156 | **0.012** |
| Recovery - Challenge | -4.278 | 0.563 | -5.378, -3.178 | -7.593 | -0.430 | **<.001** |
| Age group | 8.795 | 1.777 | 5.312, 12.274 | 4.949 | 0.376 | **<.001** |
| Challenge - Baseline x Age group | 1.097 | 1.112 | -1.074, 3.27 | 0.986 | 0.061 | 0.325 |
| Recovery - Challenge x Age group | -0.100 | 1.127 | -2.298, 2.102 | -0.089 | -0.006 | 0.929 |
| **E. Sympathetic tone** |  |  |  |  |  |  |
| Intercept | 0.109e-1 | 4.950e-4 | 1.084e-1, 1.103e-1 | 220.874 | 0.999 | **<.001** |
| Challenge - Baseline | 6.867e-4 | 5.878e-4 | -4.601e-4, 1.833e-3 | 1.168 | 0.077 | 0.244 |
| Recovery - Challenge | -1.826e-3 | 5.922e-4 | -2.983e-3, -6.719e-4 | -3.084 | -0.202 | **0.002** |
| Age group | 4.299e-3 | 9.900e-4 | 2.359e-3, 6.239e-3 | 4.343 | 0.350 | **<.001** |
| Challenge - Baseline x Age group | -1.153e-6 | 1.176e-3 | -2.295e-3, 2.291e-3 | -0.001 | 0.000 | 0.999 |
| Recovery - Challenge x Age group | -1.938e-4 | 1.184e-3 | -2.502e-3, 2.123e-3 | -0.164 | -0.011 | 0.870 |

*Note.* Models tested the fixed effects of each phase contrast of interest (Challenge - Baseline, Recovery - Challenge), age group and their interaction effects. All models included random intercepts for participants. CI = confidence interval; SE = standard error.

Examining measures of HRV during the stress task, we found that root mean square of the successive differences (RMSSD) decreased significantly from the baseline to the challenge phase (*p* = .034; Table 2) and increased significantly from the challenge to the recovery phase (*p* = .017; Table 2). Although both high-frequency (HF) and low-frequency (LF) power exhibited the same numeric pattern, the only significant phase contrast was an increase in LF power from challenge to recovery (*p* < .001; Table 2). For LF power, we also found a significant phase (challenge-baseline) x age group interaction (*p* = .012; Table 2) and a marginally significant phase (recovery-challenge) x age group interaction effect (*p* = .066; Table 2); pairwise comparisons indicated significant baseline-to-challenge decreases and challenge-to-recovery increases in LF power for younger but not older participants (Supplementary Results, Section 1).

**Table 2.** Results of linear mixed effects analyses testing that during the acute stress induction protocol, average RMSSD (A), HF power (B), and LF power (C) differed during the baseline and challenge phases and during the challenge and recovery phases.

| Predictor | Estimate | SE | 95% CI | *t* | *r* | *p* |
| --- | --- | --- | --- | --- | --- | --- |
| **A. RMSSD** |  |  |  |  |  |  |
| Intercept | 3.315 | 0.047 | 3.223, 3.407 | 70.604 | 0.987 | **<.001** |
| Challenge - Baseline | -0.066 | 0.031 | -0.127, -0.006 | -2.136 | -0.147 | **0.034** |
| Recovery - Challenge | 0.075 | 0.031 | 0.014, 0.136 | 2.412 | 0.166 | **0.017** |
| Age group | -0.556 | 0.094 | -0.74, -0.372 | -5.918 | -0.460 | **<.001** |
| Challenge - Baseline x Age group | 0.016 | 0.062 | -0.105, 0.137 | 0.257 | 0.018 | 0.797 |
| Recovery - Challenge x Age group | 0.071 | 0.062 | -0.051, 0.192 | 1.132 | 0.079 | 0.259 |
| **B. HF power** |  |  |  |  |  |  |
| Intercept | 5.869 | 0.088 | 5.696, 6.042 | 66.570 | 0.986 | **<.001** |
| Challenge - Baseline | -0.058 | 0.072 | -0.198, 0.082 | -0.811 | -0.056 | 0.418 |
| Recovery - Challenge | 0.092 | 0.071 | -0.048, 0.231 | 1.285 | 0.089 | 0.200 |
| Age group | -1.473 | 0.176 | -1.818, -1.128 | -8.354 | -0.589 | **<.001** |
| Challenge - Baseline x Age group | 0.131 | 0.143 | -0.149, 0.411 | 0.916 | 0.063 | 0.361 |
| Recovery - Challenge x Age group | 0.069 | 0.143 | -0.21, 0.347 | 0.481 | 0.033 | 0.631 |
| **C. LF power** |  |  |  |  |  |  |
| Intercept | 6.367 | 0.073 | 6.224, 6.51 | 87.135 | 0.992 | **<.001** |
| Challenge - Baseline | -0.037 | 0.067 | -0.168, 0.094 | -0.551 | -0.038 | 0.582 |
| Recovery - Challenge | 0.361 | 0.068 | 0.228, 0.493 | 5.315 | 0.347 | **<.001** |
| Age group | -1.115 | 0.146 | -1.401, -0.828 | -7.629 | -0.556 | **<.001** |
| Challenge - Baseline x Age group | 0.343 | 0.135 | 0.081, 0.606 | 2.549 | 0.173 | **0.012** |
| Recovery - Challenge x Age group | -0.251 | 0.136 | -0.516, 0.014 | -1.847 | -0.127 | 0.066 |

*Note.* Models tested the fixed effects of each phase contrast of interest (Challenge - Baseline, Recovery - Challenge), age group and their interaction effects. All models included random intercepts for participants. CI = confidence interval; LF = low-frequency; HF = high-frequency; RMSSD = root mean square of the successive differences; SE = standard error.

In terms of performance on the cognitive challenge tasks completed during the challenge phase, accuracy and reaction times on the tasks are visualized in the Supplementary Results (Section 2). On the Paced Auditory Serial Addition Task (PASAT), which was not completed by older participants, younger participants had mean accuracy of 84.3% (*SD* = 14.8%) and reaction time of 0.9 seconds (*SD* = 0.2). On the Stroop color-word matching task, younger participants were significantly more accurate than older participants, *t*(57.42) = 6.41, *p* < .001, *r* = 0.65 (*M*_younger_ = 83.7%, *SD*_younger_ = 12.2%, *M*_older_ = 62.8%, *SD*_older_ = 20.0%). Younger participants also had faster reaction times on the Stroop task compared to older participants, *t*(76.04) = -7.04, *p* < .001, *r* = -0.63 (*M*_younger_ = 1.0, *SD*_younger_ = 0.2, *M*_older_ = 1.3, *SD*_older_ = 0.2 seconds).

### 2.2. LC MRI contrast in the sample

Peak, rostral and caudal LC contrast values in the sample are presented in Figure 2. A Welch’s *t*-test indicated a trend toward higher peak LC contrast in older relative to younger participants, *t*(114.47) = 1.88, *p* = .063, *r* = 0.173. A 2x2 mixed analysis of variance to examine effects of topography (rostral/caudal) and age group indicated a significant main effect of age group on LC contrast, *F*(1, 151) = 4.05, *p* = .046, *r* = 0.162, driven by higher ratios in older relative to younger participants, but no significant effect of topography, *F*(1, 151) = 1.16, *p* =.283, *r* = 0.087, or age group x topography interaction on LC contrast, *F*(1, 151) = 2.48, *p* = .117, *r* = 0.127.

**Figure 2.**
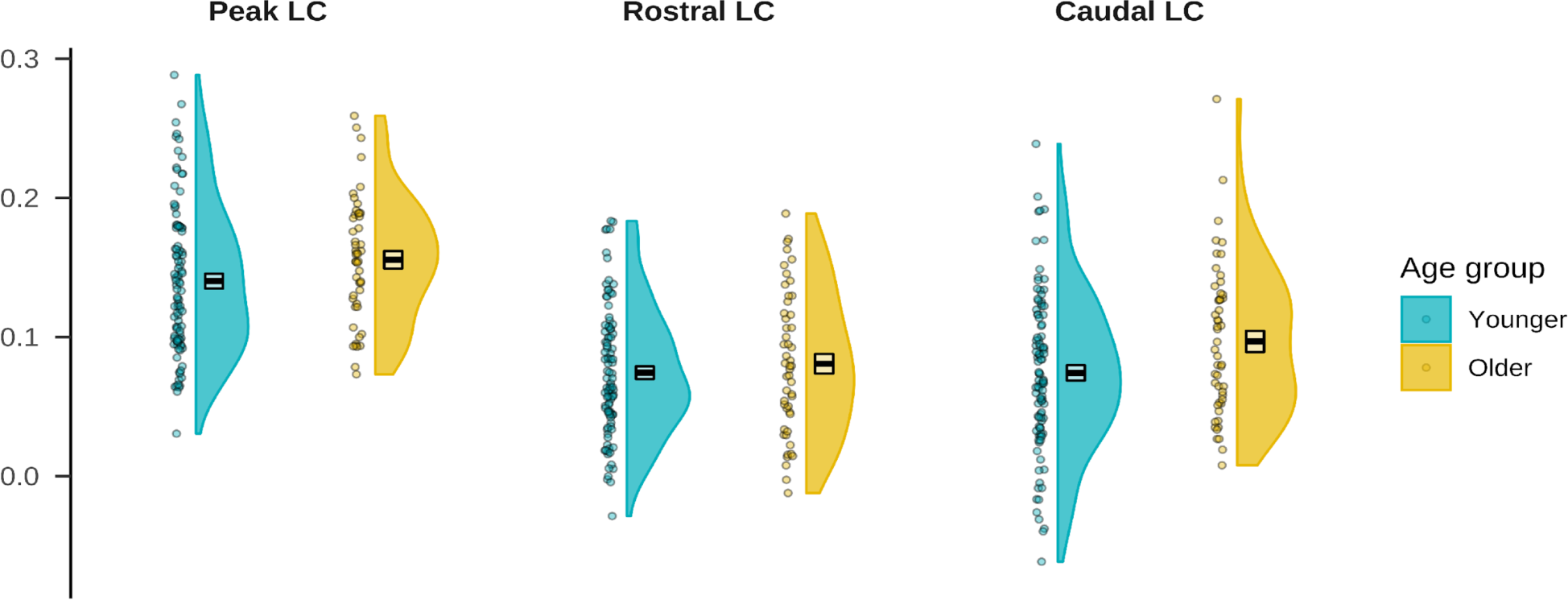
LC MRI contrast in the sample.

### 2.3. Associations between LC MRI contrast and arousal during the stress induction task

After calculating values of acute stress reactivity and recovery for each arousal measure, pairwise Pearson correlation analyses were used to assess associations between LC MRI contrast and arousal during rest, stress reactivity and stress recovery. Visualizations of pairwise correlations between LC contrast values and all arousal measures are presented in Figure 3. Pearson’s correlation analyses indicated that in younger participants, peak LC contrast was significantly negatively correlated with breathing rate during stress recovery, *r*(95) = -0.21, *p* = .038, and rostral LC contrast was positively correlated with sympathetic tone during stress recovery, *r*(64) = 0.31, *p* = .010. In older participants, peak LC contrast was significantly positively correlated with LF power at baseline, *r*(41) = 0.31, *p* = .045. There was also a marginally significant negative correlation between caudal LC contrast and HF power during stress recovery in older participants, *r*(33) = -0.32, *p* = .058. No other arousal measures were significantly correlated with LC contrast.

**Figure 3.**
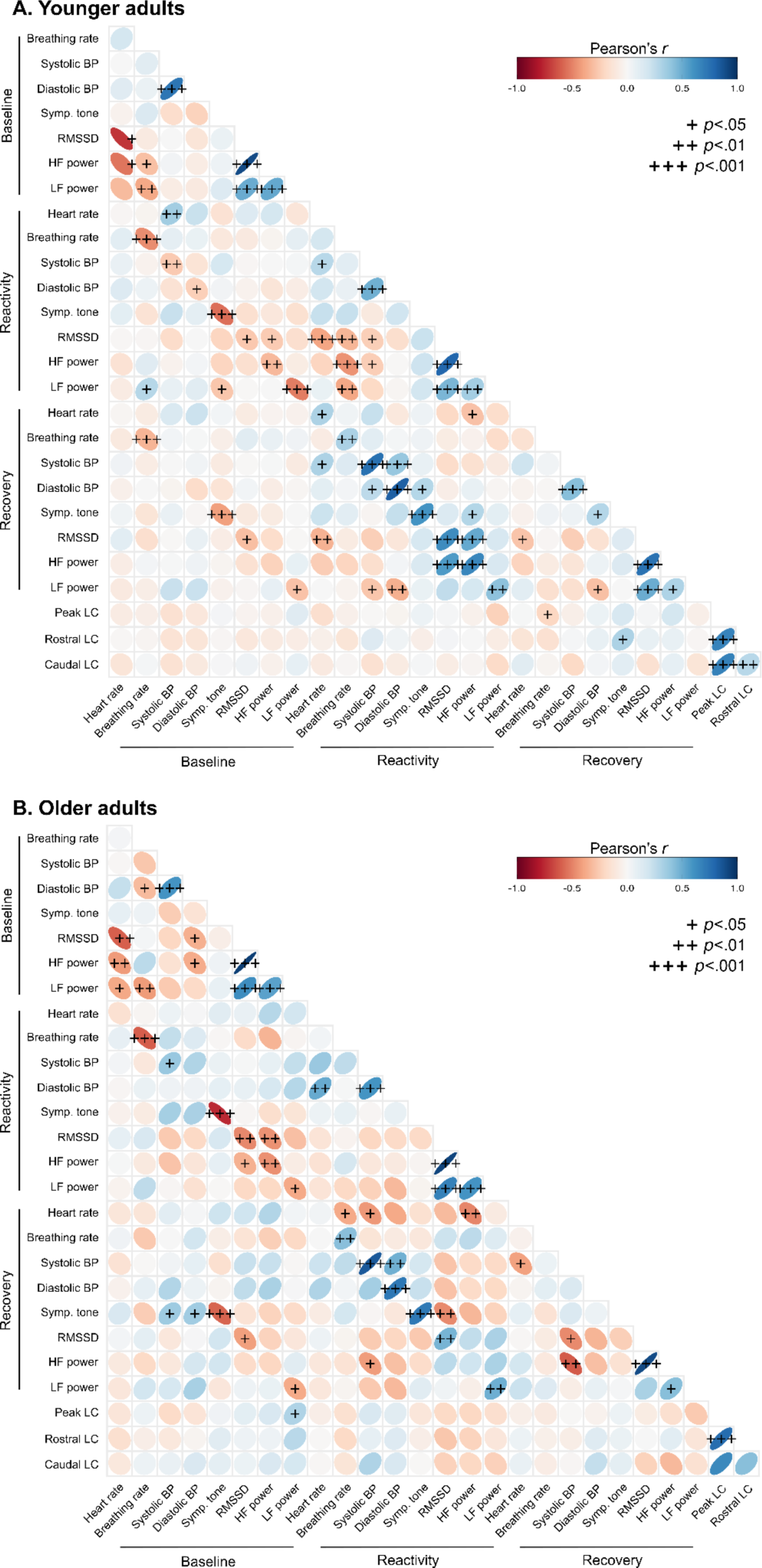
Visualization of correlation matrices reflecting pairwise Pearson correlations between LC contrast and arousal during the stress induction protocol. + *p*<.05, ++ *p*<.01, +++ *p*<.001.

As a multivariate approach to quantify the associations between LC contrast and arousal during the stress induction protocol, we performed a series of partial least squares (PLS) correlation analyses. We note that these analyses were performed with only participants with no missing values (53 younger, 23 older), whereas the correlations presented above reflected all available sets of pairwise observations. PLS analyses examining associations of peak and rostral LC contrast, respectively, with arousal yielded no reliable latent variables. The final PLS analysis, examining associations between caudal LC contrast and arousal, indicated 1 marginally reliable latent variable (*p* = .058). Bootstrap ratios reflecting the contribution of each arousal measure to this latent variable, as well as the correlation between physiological scores for this latent variable and peak LC contrast values, are shown in Figure 4. Physiological scores on this latent variable were highly correlated with caudal LC contrast in older participants, *r*(21) = 0.63, *p* = .001, but not in younger participants, *r*(51) = 0.10, *p* = .487. Furthermore, higher physiological scores for this latent variable reflected greater systolic blood pressure increases and RMSSD decreases during stress reactivity, higher systolic blood pressure during stress recovery, and lower RMSSD and HF power during stress recovery.

**Figure 4.**
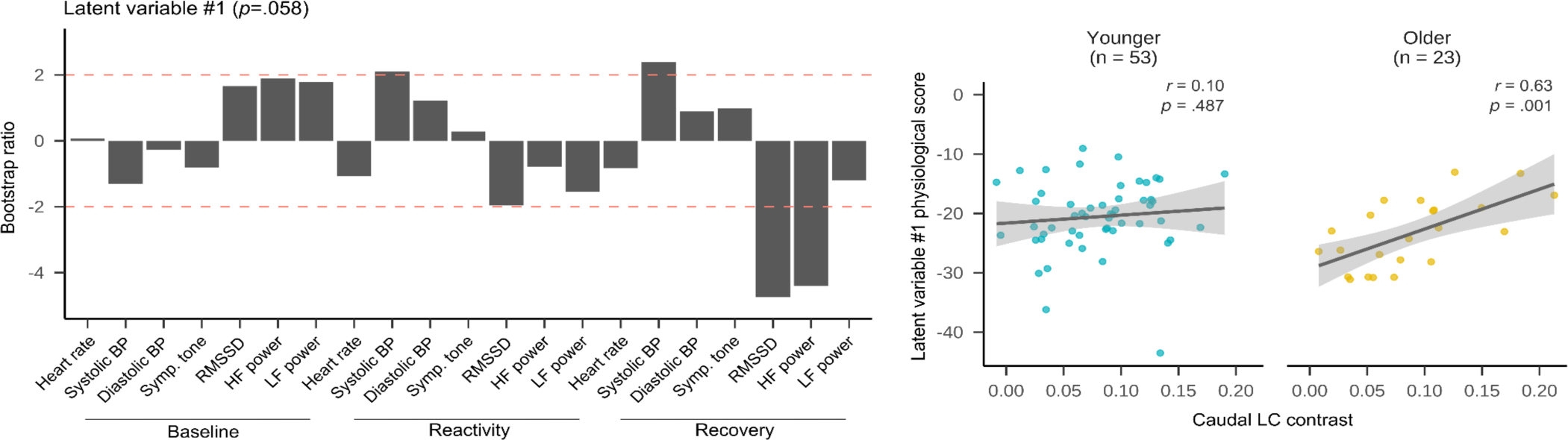
Results of partial least squares (PLS) correlation analyses examining the association between caudal LC contrast and physiological arousal during the stress induction task. These analyses indicated a marginally reliable latent variable reflecting an association between caudal LC contrast and arousal for older participants. The left panels depict bootstrap ratios which reflect how much each arousal measure contributed to the latent variable (bootstrap ratios with absolute value greater than 2, indicated in red, were considered stable contributors). Right panels depict associations between physiological scores - reflecting the projection of each respective latent variable onto the original physiological arousal data - and LC contrast values.

## 3. Discussion

As an arousal hub region in the brain, the LC plays a major role in the central nervous system’s response to acute stress, releasing norepinephrine throughout the brain and spinal cord to promote behaviors that facilitate stressor avoidance or elimination (Bremner et al., 1996; Sara & Bouret, 2012; Wood & Valentino, 2017). Studies using MRI to assess the LC’s structure in vivo have suggested that having a more structurally intact LC in later adulthood is associated with better cognitive outcomes and reduced risk of cognitive decline (Dahl et al., 2019; Elman et al., 2021a; Liu et al., 2020), but it is unclear whether LC MRI contrast is related to acute stress responses. Here, we tested how LC contrast was associated with physiological responses during rest, reactivity to acute stress, and recovery from acute stress in both younger and older adults. Across univariate and multivariate analyses, we found that for older adults, having higher caudal LC contrast was associated with higher stress-related increases in systolic blood pressure and lower HRV during stress recovery. Together, these findings suggest that having higher caudal LC contrast in older adulthood is associated with more pronounced physiological responses to acute stress.

In response to acute psychosocial stressors, sympathetic arousal increases and HRV generally decreases, reflecting parasympathetic withdrawal (Rab & Admon, 2021). Aging is associated with changes to the autonomic system that may impact acute stress responses (Kaye & Esler, 2008). In general, cortisol levels are higher in older adults than in younger adults, reflecting higher tonic activation of the HPA axis in later adulthood (Lupien et al., 2009). Sympathetic nervous system activity tends to also increase in aging (Fagius & Wallin, 1993; Seals & Esler, 2000), and consistent with this pattern, we found that older participants had higher values of a measure of skin sympathetic nerve activity quantified from ECG signals, relative to younger participants. Vagal control of heart rate and HRV also typically decline with age (Jandackova et al., 2016), and consistent with this pattern, we found that older participants had lower HRV values than younger adults. Evidence for age changes in parasympathetic responses to stress is scarce, but here, we found that older participants had significantly smaller decreases in LF power relative to younger participants during acute stress. Thus an autonomic system that is relatively less affected by age-related dysregulation might be expected to feature more dynamic responses to acute stress - specifically, greater sympathetic increases and greater parasympathetic withdrawal in response to stress.

We found that older participants with more dynamic physiological responses to acute stress - that is, greater sympathetic increases and parasympathetic withdrawal in response to stress - had higher caudal LC contrast. These findings add to a growing body of literature linking higher LC MRI contrast to better cognitive and neural outcomes in aging (Bachman et al., 2021; Dahl et al, 2019; Elman et al., 2021b; Liu et al., 2020), but they also support the notion that the LC is important for allowing messages from the brain to reach the heart to coordinate effective autonomic responses in aging. Indeed, as a component of the central autonomic network (Benarroch, 1993), the LC contains excitatory projections to the rostral ventrolateral medulla, inhibitory projections to parasympathetic nuclei, and bidirectional connections with C1 neurons that coordinate cardiovascular responses (Lamotte et al., 2021).

Our results furthermore highlight a potential role of the LC’s caudal aspect in neurovisceral integration. We previously found that in aging, associations with episodic memory and gray matter integrity were greater for rostral than caudal LC contrast (Bachman et al., 2021; Dahl et al., 2019). Consistent with the rostral LC being important for age-related outcomes, the rostral LC undergoes relatively more cell loss than the caudal LC in aging (Manaye et al., 1995) and Alzheimer’s disease (Zarow et al., 2003). So why might the caudal LC be more relevant for physiological arousal in aging? Hirschberg et al. (2017) identified two populations of LC neurons: one population consisting of more rostrally-originating neurons projecting to the prefrontal cortex and another clustered in the caudal LC and projecting primarily to the spinal cord. At the spinal cord, noradrenergic neurons from the LC synapse onto sympathetic preganglionic neurons, promoting downstream peripheral arousal responses (Clark & Proudfit, 1991). Based on current evidence, it is unclear whether medullary and parasympathetic projections from the LC also originate predominantly in the LC’s caudal aspect. Yet our results support the possibility that the caudal LC plays a role in the pathway linking neural appraisals of the world to cardiovascular responses.

We found associations between LC contrast and patterns of acute stress responding in older participants, but we did not find the expected relationships between LC contrast and arousal in younger participants. Although a limited number of studies have investigated associations with LC structure in younger adults, the largely null findings in our younger sample are inconsistent with reports of higher LC volume being associated with higher anxious arousal (Morris et al., 2020b) and of caudal LC contrast being negatively associated with cortical thickness (Bachman et al., 2021) in younger adults. We previously reported that LC contrast was negatively correlated with HRV during a fear conditioning task in both younger and older adults (Mather et al., 2017). Our current findings offer an alternative explanation for the previous findings in older adults (more pronounced acute stress responses in individuals with higher LC contrast), but they are inconsistent with the previous findings in younger adults. In this case, a potential reason that we did not observe expected associations in younger adults is that the LC contrast measure does not reflect functional acute stress responses in younger adults. Furthermore, if LC contrast reflects neuromelanin, the imaging method used may be less reliable in younger than older adults due to relatively less neuromelanin accumulation in LC neurons (Betts et al., 2019b; Manaye et al., 2005).

Another potential explanation for the lack of findings in younger participants is that the challenge tasks may not have reliably induced acute stress for the younger cohort. Qualitatively, older participants reported the Stroop task to be very challenging, whereas this was not common feedback from younger participants. Heart rate, breathing rate, systolic and diastolic blood pressure and sympathetic tone all increased reliably for younger participants during the challenge phase. However, because we did not measure salivary cortisol levels throughout the acute stress induction task, we cannot be sure that this task elicited an acute stress response in participants or simply cognitive load. In addition, a limitation of the study is that the stress induction protocol contained 2 tasks for younger participants and 1 task for older participants. We re-ran all analyses using only data from the Stroop task to calculate stress reactivity for all participants, but this did not affect the pattern of results. Further research utilizing equivalent methods of stress induction in younger and older adults will be important for confirming our findings.

We used pairwise correlation analyses to assess relationships between LC contrast and physiological stress responses, but we note that the number of statistical tests performed is a limitation of this approach. To combat this limitation, and as a multivariate alternative to assess relationships with LC contrast, we used a multivariate method, PLS correlation. Yet PLS also suffers from an important limitation: Specifically, for this method, we used only cases with all available physiological measures, constraining the number of participants included for these analyses. The PLS results suggest the interesting possibility that older adults with caudal LC contrast have more pronounced physiological responses to acute stress, but further work is needed to replicate these findings in a larger sample of older adults. We also note a final limitation of these analyses: HF power was a reliable contributor to the latent variable identified by PLS, but respiration can influence HF and LF power (Laborde et al., 2017). Because the challenge tasks entailed significant changes in breathing rates, the HF and LF power results should be interpreted with caution.

To conclude, we examined how LC MRI contrast was related to physiological arousal at rest, during acute stress reactivity, and during recovery from acute stress. In younger participants, LC contrast was largely unrelated to physiological responses to stress, although this may be explained by the stress induction task being challenging, but not reliably stressful, for younger participants. In older participants, caudal LC contrast was associated with greater stress-related increases in systolic blood pressure and decreases in HRV, as well as lower HRV during stress recovery. These results support the novel hypothesis that caudal LC structure is associated with more pronounced stress responses in aging.

## 4. Methods and materials

### 4.1. Participants

Data were collected as part of a clinical trial testing the effects of heart rate variability biofeedback training on emotion regulation brain networks (Nashiro et al., 2021). For the present analyses, only data from the pre-intervention measurement timepoint - that is, before participants learned about or started the intervention - were considered. Specifically, we considered data from all participants who completed an MRI session including a TSE scan and an acute stress induction task at the pre-intervention timepoint. This included 115 younger and 59 older participants (data collection for the older cohort was terminated prematurely due to the COVID-19 pandemic). These participants were MRI-eligible individuals without major medical, neurological, psychiatric, or cardiac conditions. Individuals who engaged in regular relaxation, biofeedback or breathing techniques were excluded from participation, as were individuals taking psychoactive medications. Individuals taking antidepressants or anti-anxiety medications were eligible to participate so long as the medication had been taken for at least three months prior to study participation. Older adults who scored lower than 16 on the TELE, a brief cognitive assessment administered over the telephone (Gatz et al., 1995), were excluded from participation for possible dementia.

Of those who completed the acute stress induction task and an MRI session, scans from 18 participants were excluded from LC delineation due to severe motion artifact (n = 15), susceptibility artifact overlapping the LC or pons (n = 2), and incorrect scan resolution (n = 1). Following LC delineation, 1 older participant was excluded from analysis due to incorrect placement of the LC search space, and 2 participants (1 younger, 1 older) were missing complete physiological recordings from the stress task due to recording errors and were therefore excluded from analysis. The final sample for analysis included 153 participants (102 younger, 51 older) and is described in Table 3. The study protocol was approved by the University of Southern California Institutional Review Board. All participants provided written, informed consent and received monetary compensation for their participation.

**Table 3.** Sample characteristics.

|  | Younger (N = 102) | Older (N = 51) | Comparison,<br>*p*-value |
| --- | --- | --- | --- |
| Age, mean (SD) | 22.66 (2.89) | 64.61 (6.31) | <.001 |
| Age, range | 18 - 31 | 55 - 80 |  |
| N (%) females | 55 (53.9%) | 34 (66.7%) | 0.183 |
| Education, mean (SD) | 15.9 (2.16) | 16.51 (2.44) | 0.131 |
| Education, range | 12 - 24 | 12 - 25 |  |
| CESD, mean (SD) | 13.95 (7.54) | 9.93 (7.09) | 0.002 |
| CESD, range | 1.5 - 39.5 | 0 - 28.5 |  |
| TAI, mean (SD) | 42.15 (9.97) | 35.3 (10.54) | <.001 |
| TAI, range | 21.5 - 65.5 | 20 - 67 |  |

*Note.* Age and education are presented in years. During their lab visits for the acute stress induction protocol and MRI session, participants completed the Center for Epidemiological Studies Depression Scale (CES-D; Radloff, 1977) and Trait Anxiety Inventory (TAI; Spielberger et al., 1983). A single score for each participant was calculated by averaging over their scores from both visits. Comparison *p*-values reflect results of independent samples Welch's *t*-tests and, for comparing the proportion of females in each age group, a chi-squared test.

### 4.2. Data collection

#### 4.2.1. Acute stress induction protocol

During the first week of the study, participants completed a computerized acute stress induction procedure based on a standardized protocol known to elicit a robust acute physiological stress response (Crowley et al., 2011). The task consisted of a 4-minute baseline resting phase, a cognitive challenge phase, and a 4-minute recovery resting phase (Figure 5).

**Figure 5.**
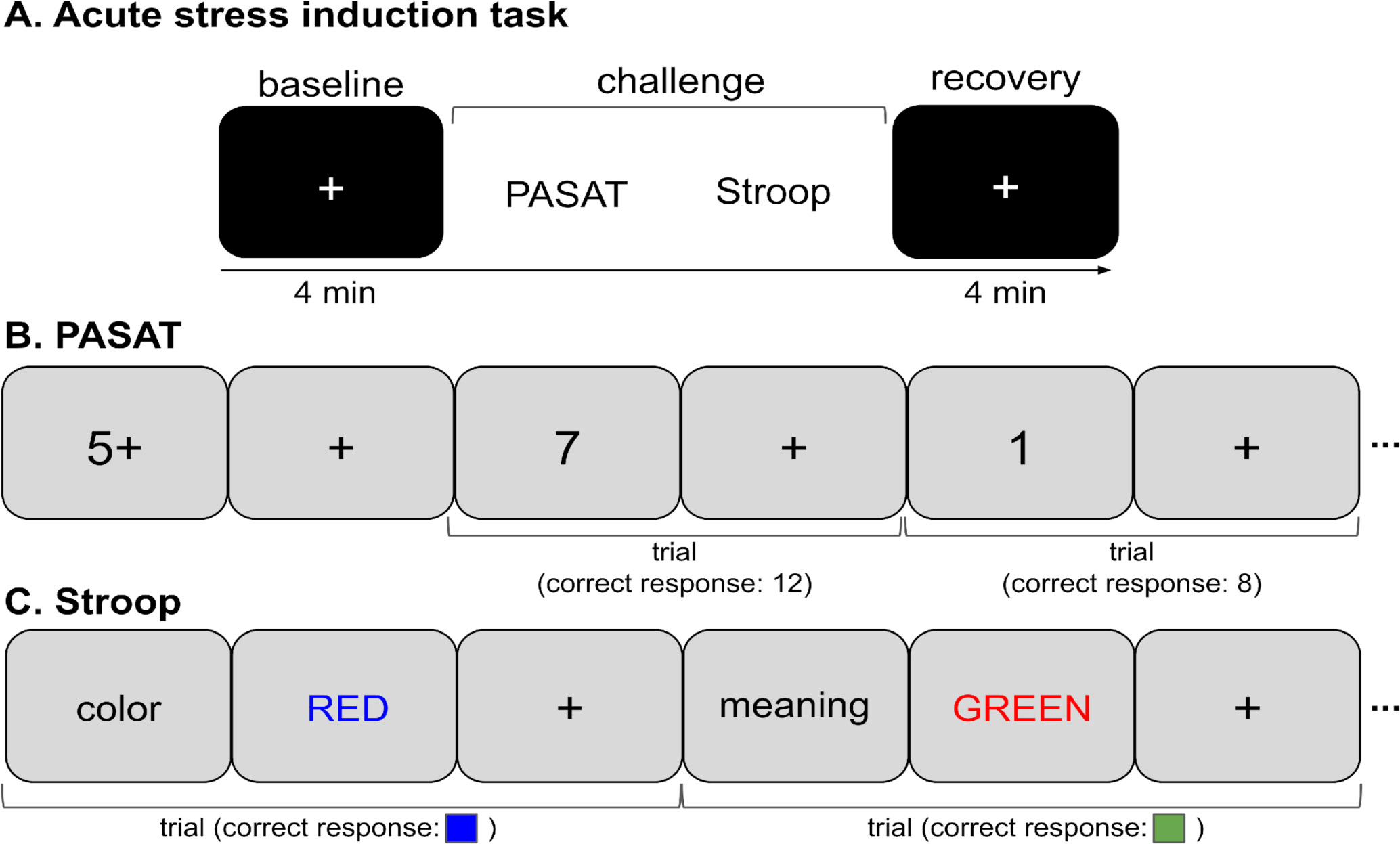
Acute stress induction protocol. The protocol consisted of 3 phases (A): a baseline resting phase, a cognitive challenge phase, and a recovery resting phase. The challenge phase consisted of two tasks: the Paced Auditory Serial Addition Task (PASAT; B) and a Stroop color-word matching task (C). Younger adults completed both tasks, whereas older adults completed only the Stroop task.

During the baseline phase, participants sat with their feet resting flat on the ground and their hands resting in a supine position on a flat tabletop. Participants viewed a black screen with a white fixation cross centered on the screen and were instructed to refrain from talking and to breathe normally during this phase. Following the baseline phase, participants completed a challenge phase which consisted of two cognitive tasks for younger adults and one task for older adults (piloting indicated that the first task was excessively frustrating for older participants). To increase the socially evaluative nature of the challenge phase, participants were told that their performance would be evaluated by the experimenter and compared with that of other participants. The first task, completed only by younger adults, was a computerized version of the Paced Auditory Serial Addition Task (PASAT; Figure 5B; Tombaugh, 2006) in which participants were presented with a series of digits and instructed to add each digit to the digit that came directly before it. Participants were instructed to enter the resulting sum on the keyboard using their dominant hand, and potential responses were never greater than 20. The task consisted of 30 trials in which participants had 3 seconds to respond to each digit; the task lasted approximately 160 seconds.

The second task, completed by both younger and older adults, was a Stroop color-word matching task (Figure 5C; MacLeod, 1991), in which a color word (‘RED‘, ‘BLU‘, or‘GREEN‘) was presented on a computer screen in a color incongruent with its meaning (either red, green or blue). Participants were instructed to use their dominant hand to press a key corresponding to either the color in which the word was presented, or the meaning of the word, based on an instruction which appeared directly before the word. The Stroop task consisted of 20 trials and lasted approximately 120 seconds.

During both tasks, auditory feedback was provided to participants on each trial: A bell sound was played in response to correct responses, whereas a buzzer sound was played after missing or incorrect responses to increase the socially-evaluative nature of the tasks (Dickerson & Kemeny, 2004). Before beginning the tasks, participants were provided with instructions and practice trials for both tasks. Following the cognitive challenge phase, participants completed a 4-minute recovery resting phase that was identical to the baseline resting phase.

Physiological signals were recorded throughout the acute stress induction protocol at a sampling rate of 2KHz using a BIOPAC MP160 system (Goleta, CA). ECG signals were collected with a standard Lead II configuration with disposable, pre-gelled Ag/AgCl electrodes (EL501) and transmitted using a wireless BioNomadix transmitter system. Respiration was measured with the Biopac Respiratory Effort Transducer, which involved a belt being placed around the lower rib cage to measure changes in chest circumference, and signals were transmitted using the BioNomadix system. Continuous blood pressure was recorded on each participant’s non-dominant arm using a BIOPAC noninvasive blood pressure monitoring system (NIBP100D).

#### 4.2.2. Magnetic resonance imaging

Approximately one week after completing the stress induction task, participants returned to the laboratory for an MRI session. Sequences of interest for the present analyses included a three-dimensional, T1-weighted magnetization prepared rapid gradient echo (MPRAGE) anatomical scan and a TSE scan covering the entire pons. Parameters of the MPRAGE sequence were as follows: TR = 2300 ms, TE = 2.26 ms, flip angle = 9°, field of view = 256 mm, voxel size = 1.0 x 1.0 x 1.0 mm, 175 volumes collected. Based on the MPRAGE scan, a two-dimensional, multi-slice TSE scan covering the entire pons was collected. The TSE sequence had the following parameters: TR = 750 ms, TE = 12 ms, flip angle = 120°, bandwidth = 287 Hz/pixel, voxel size = 0.43 x 0.43 x 2.5 mm, gap between slices = 1.0mm, 11 axial slices.

### 4.3. Physiological data analysis

Physiological signals collected during the acute stress induction protocol were first split into segments for each participant, with segments corresponding to the various parts of the protocol: baseline, PASAT, Stroop task, and recovery phase. Quality control checks and preprocessing were performed for each segment separately, using the steps described below. For 5 older participants, we included only baseline segments for preprocessing and analysis, because these participants completed a pilot version of the protocol that included 2 cognitive challenge tasks. Processing steps described in this section were performed using MATLAB (Version R2021b).

#### 4.3.1. Preprocessing and quality control

##### ECG signals

To remove baseline wander and high-frequency noise, ECG segments were filtered with a finite impulse response (FIR) bandpass filter with a passband between 0.5 and 40 Hz. ECG signals were assessed for quality in two steps. ECG segments were first visually inspected for signal quality and noise; 12.3% (n = 68) of all segments demonstrating excessive noise or abnormalities such that QRS-complexes were not discernible were excluded from analyses of heart rate and HRV. Second, during r-peak delineation and HRV analysis (see Section 4.3.2), an average signal quality index from 0-1 reflecting a comparison between r-peak annotations performed by two algorithms, jqrs and wqrs (Behar et al., 2014; Johnson et al., 2014), was calculated for each segment. We excluded an additional 5.0% (n = 24) of ECG segments for having an average signal quality index of below 0.7. Further segments were excluded for atrial fibrillation being detected (6.4%, n = 31) and having more than 20% of peaks missing (0.8%, n = 4).

##### Respiration signals

Respiration segments were resampled to 50 Hz and filtered with an FIR filter with a passband between 0.05 and 1 Hz. Preprocessed segments were then visually inspected for signal quality and were overlaid with detected peaks corresponding to inhalations, for visual inspection of peak detection accuracy. Respiration segments with poor quality and/or inaccurate peak detection in the majority of the segment were excluded from all analyses. This led to 6.4% (n = 35) of respiration segments being excluded from analysis of breathing rate.

##### Continuous blood pressure signals

For removal of high-frequency noise from continuous arterial blood pressure segments, we applied a FIR lowpass filter with a cutoff frequency of 40 Hz. Raw segments were visually inspected for abnormalities; those in which regular systolic peaks were not detectable were excluded from analyses of systolic blood pressure. This led to 3.6% (n = 20) of segments being excluded from analysis. For another 37 segments, the blood pressure monitor re-calibrated mid-way through data collection; these segments were also excluded from analysis.

#### 4.3.2. Calculation of arousal measures

Preprocessed physiological data segments were next used to compute segment-wise measures of physiological arousal. We calculated average values of each arousal measure across the baseline, challenge and recovery phases for each participant. For younger participants, a single value of each measure for the challenge phase was calculated by averaging values from the Stroop and PASAT segments^1^. The resulting average values were first used for visualization and for testing whether the acute stress induction task effectively modulated arousal in each age group (see Section 4.5.2). We then used the average values to calculate stress reactivity and recovery, as described in Section 4.5.3. Calculation of arousal measures was performed using MATLAB (Version R2021b).

##### Heart rate and heart rate variability

The PhysioNet Cardiovascular Signal Toolbox (Version 1.0.2; Goldberger et al., 2000; Vest et al., 2018), an open-source toolbox designed to address issues of validation, standardization and reproducibility in HRV signal processing, was used for QRS-complex detection and to calculate time- and frequency-domain measures of HRV from preprocessed ECG signals. QRS detection was performed with the jqrs beat detector (Behar et al., 2014; Johnson et al., 2014). Parameters for HRV calculation are described in the Supplementary Methods (Section 1). Based on delineated r-peaks, we calculated a value of mean heart rate for each segment.

Time-domain HRV analysis was performed on resulting rr-intervals in each segment, yielding a measure of RMSSD for each segment. Frequency-domain analysis was performed for each full segment using the Lomb periodogram method for generating power spectral density, which is the default in PhysioNet because it can handle losses of up to 20% of data (Clifford, 2002). This yielded measures of LF and HF spectral power for each segment. Frequency bands of 0.04 - 0.15 and 0.15 - 0.4 Hz were used for calculating LF and HF power, respectively.

##### Breathing rate

Peaks corresponding to inhalations were identified on resampled, filtered respiration signals using MATLAB’s ‘findpeaks‘ function with a minimum peak width of 500 milliseconds. Identified peaks were used to calculate a single value of breathing rate for each segment.

##### Systolic and diastolic blood pressure

Beat-to-beat systolic and diastolic blood pressure were extracted from preprocessed continuous blood pressure segments using the algorithm from the CRSIDLab toolbox (da Silva & Oliveira, 2020). This method uses rr-intervals in the corresponding ECG segment to identify systolic peaks and dicrotic notches within each cardiac cycle (Parati et al., 1995). For this method, a systolic/diastolic threshold of 80 was specified (the default in CRSIDLab). When rr-intervals were unavailable due to a low-quality ECG segment, an alternative algorithm based on the continuous blood pressure waveform was used (Li et al., 2010). For both methods, we specified that successive maxima and successive minima could be a minimum of 0.375 and a maximum of 2 seconds apart. Identified peaks were used to calculate a value of mean systolic and diastolic blood pressure for each segment.

##### Sympathetic tone

We assessed sympathetic tone using the neuECG method, an approach to quantify skin sympathetic nerve activity from ECG signals (Kusayama et al., 2020). In this approach, raw ECG segments were first high-pass filtered with an FIR filter with cutoff frequency of 500 Hz. Filtered ECG signals were then full-wave rectified and integrated with a leaky integrator (time constant: 0.1 seconds). We then computed the average voltage of the resulting signal across each segment (aSKNA), a measure that has been shown to increase during sympathetic-activating manipulations (Kusayama et al., 2020).

### 4.4. Locus coeruleus delineation

We used a validated, semi-automated approach to delineate the LC on TSE scans (Figure 6). The method is fully described in Bachman et al. (2021) and based on approaches by Dahl et al. (2019) and Ye et al. (2021). As a summary, LC delineation entailed aligning all MPRAGE and TSE scans across participants, and then warping TSE scans to MNI152 linear 0.5mm standard space. LC delineation steps were performed using Advanced Normalization Tools (ANTs, Version 2.3.4, Avants et al., 2011), and visualization steps were performed using ITK-SNAP (Version 3.6.0, Yushkevich et al., 2006).

**Figure 6.**
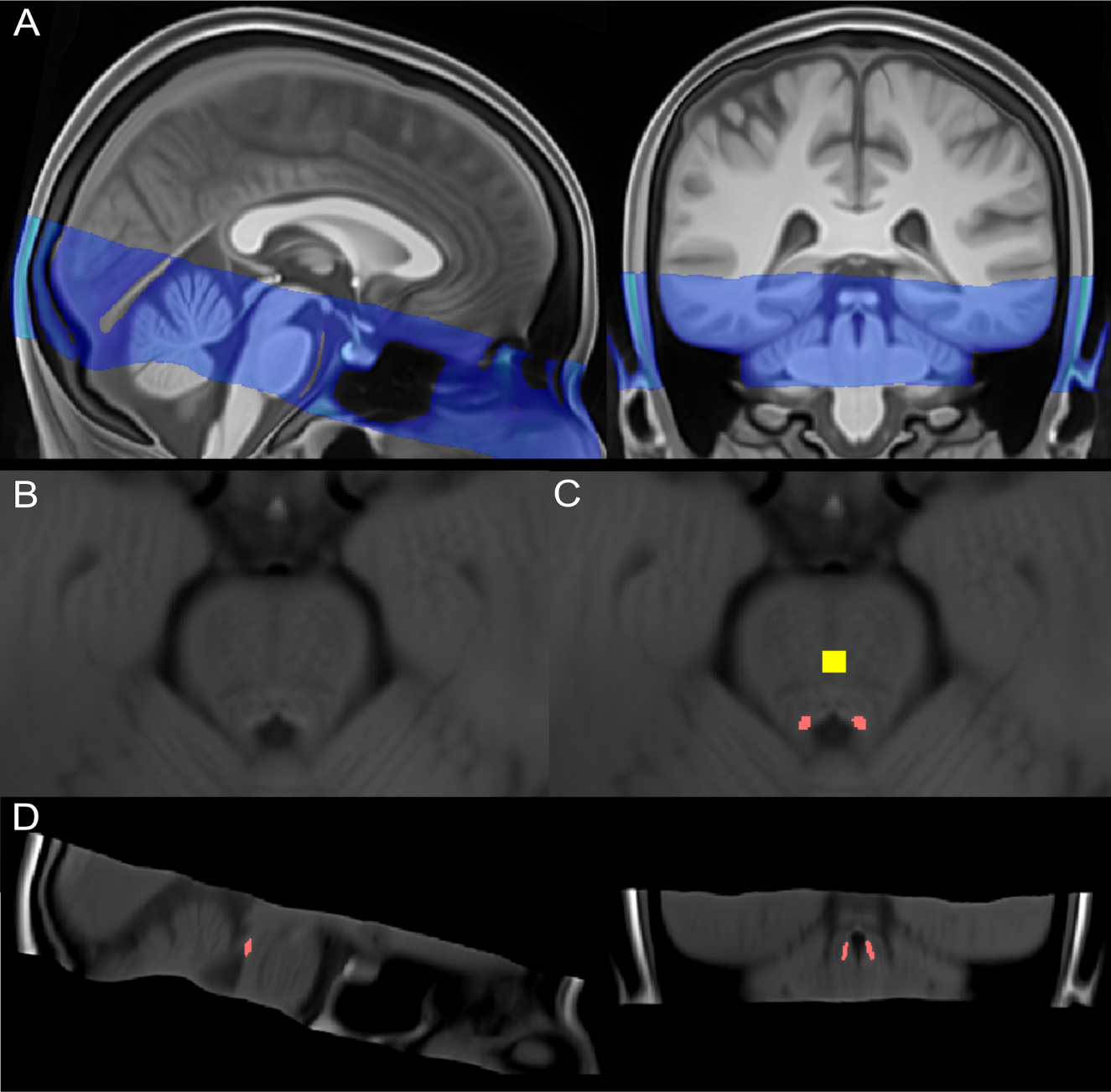
LC delineation procedure. (A) TSE template overlaid on the MPRAGE template, both warped to MNI152 (linear) 0.5mm standard space. (B) Detailed view of pons on TSE template in standard space, including hyperintensities bordering the 4th ventricle. (C) LC map from Dahl et al. (2022) shown in pink, overlaid on the TSE template in standard space. Reference map, also from Dahl et al. (2022), shown in yellow. (D) Sagittal (left) and coronal (right) views of LC map (pink) overlaid on the TSE template in standard space.

To isolate potential LC voxels on warped TSE scans, we applied the Dahl et al. (2022) meta-map as a mask; this mask was selected because it aggregates across many published LC maps. Within the masked region on each scan, we extracted the intensity value and coordinates of the peak-intensity voxel in each slice in the z dimension for left and right LC. Locations of peak LC intensity are depicted in the Supplementary Methods (Section 2). As a validation step, two raters delineated the LC on native TSE scans (see Supplementary Methods, Section 3). LC intensities determined through the semi-automated approach were found to have high correspondence between LC intensities from manual delineation, based on intra-class correlation coefficients calculated using consistency two-way models (left LC: *ICC*(3,1) = 0.948, 95% CI = 0.929 - 0.962, *p* < .001; right LC: *ICC*(3,1) = 0.938, 95% CI = 0.915 - 0.955, *p* < .001). As a reference region for calculating LC contrast ratios, we also applied the Dahl et al. (2022) central pontine reference map as a mask on each warped TSE scan and extracted the peak intensity within the masked region in each z-slice. Values of left and right contrast for each participant and z-slice were then calculated as a ratio (Liu et al., 2017):

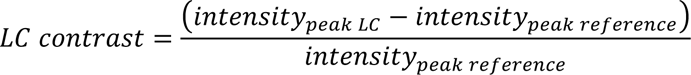

Contrast ratios for left and right LC were averaged within each z-slice, and in a final step, the peak ratio across all z-slices was used for statistical analysis (these are henceforth referred to as peak LC contrast values). Based on evidence that rostral LC exhibits greater neuronal loss in aging and Alzheimer’s disease relative to caudal LC (Manaye et al. 1995; Zarow et al., 2003), as well as reports of spatially confined associations with contrast along the LC’s rostrocaudal axis (Bachman et al., 2021; Dahl et al., 2019), we also calculated rostral and caudal LC contrast values for each participant. This entailed first calculating percentiles of slices along the LC’s rostrocaudal axis where we previously found age differences in LC contrast and associations with cortical thickness in younger versus older adults (Bachman et al., 2021). These percentiles were applied to the z-range of slices included in the LC meta-map (*z* = 85-112) to identify ranges of slices corresponding to rostral and caudal LC (rostral: MNI *z* = 101-104; caudal: MNI *z* = 87-95). For each participant, contrast ratios were then averaged across the caudal and rostral clusters of slices to obtain values of rostral and caudal LC contrast, respectively.

### 4.5. Statistical analysis

#### 4.5.1. Data transformation and outlier removal

RMSSD, LF power and HF power values were determined to exhibit severe non-normality and were therefore log-transformed prior to analysis. Outliers for each arousal measure were then identified using the mean absolute deviation-median rule for each age group separately and treated as missing values for all analyses (Wilcox, 2011). Outlier detection was performed prior to the detection of peak (or minima) arousal metrics for each challenge segment. A summary of identified outliers for each measure is included in the Supplementary Methods (Section 4).

For peak, rostral and caudal LC contrast values, we tested for outliers for younger and older adults separately according to the mean absolute deviation-median rule. One older participant was an outlier for peak LC ratios and excluded from relevant analyses.

#### 4.5.2. Analysis of effectiveness of the acute stress induction protocol

To assess whether the acute stress induction protocol was effective at modulating each measure of physiological arousal in younger and older adults, we fit a series of linear mixed effects models. These models tested the fixed effects of phase (baseline/challenge/recovery), age group, and their interaction on each arousal measure (heart rate, breathing rate, systolic blood pressure, diastolic blood pressure, sympathetic tone, RMSSD, LF power and HF power), using a separate model for each measure. For these analyses, we used average values of each measure from each phase. We used a repeated contrast coding scheme for the phase factor to test two contrasts of interest: challenge vs. baseline and recovery vs. challenge (Schad et al., 2020). Age group was sum coded (younger = -0.5, older = 0.5). Models were fit with the ‘lmer4‘ R package (Version 1.1-27.1; Bates et al., 2015) and parameter significance was assessed with the ‘lmerTest‘ package using Satterthwaite’s method (Version 3.1-3; Kuznetsova et al., 2017). Planned comparisons of each measure for each phase contrast were performed with Bonferroni corrections for multiple comparisons, using the ‘emmeans‘ package (Version 1.7.0; Lenth, 2021).

We also examined performance on the PASAT and Stroop tasks by computing each participant’s mean accuracy and reaction time on the tasks. As only younger adults completed the PASAT, we reported PASAT performance as the mean and standard deviation of accuracy and reaction time across participants. For the Stroop task, we used independent-samples Welch’s t-tests to compare accuracy and reaction times by age group.

#### 4.5.3. Calculation of measures of acute stress reactivity and recovery

We then calculated measures of acute stress reactivity by computing the change in each physiological measure from baseline to the challenge phase (Llabre et al., 1991). For these analyses, we used average values of each measure from each phase:

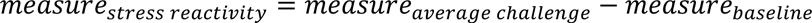

To calculate measures of acute stress during recovery, we likewise computed the difference in each measure from baseline to the recovery phase:

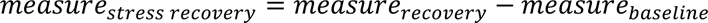

Larger-magnitude values of reactivity were therefore expected to reflect greater stress reactivity, whereas larger-magnitude values of recovery would reflect higher arousal during stress recovery.

#### 4.5.4. Analysis of LC MRI contrast in the sample

Prior to testing associations between LC contrast and arousal, we examined whether peak LC contrast differed in younger and older adults using an independent-samples Welch’s *t*-test. Based on previous findings of age differences in contrast according to LC topography (Dahl et al., 2019), we also performed a separate 2x2 mixed-design analysis of variance, implemented with the R package ‘afex‘ (Version 1.0-1; Singmann et al., 2021), testing the effects of age group (younger, older) and topography (rostral, caudal) on LC contrast.

#### 4.5.5. Analysis of associations between LC MRI contrast and physiological arousal

We first assessed associations between LC contrast and physiological arousal by computing, separately for younger and older participants, a Pearson correlation matrix reflecting pairwise correlations between all measures of arousal (baseline, reactivity and recovery) and all measures of LC contrast (peak, rostral and caudal). For this step, all available pairwise observations were used.

To further probe associations between LC contrast and arousal using a multivariate framework, we then used a series of PLS correlation analyses. The aim of PLS is to identify latent variables that express a maximal amount of covariance between a set of predictors and an outcome variable (Krishnan et al., 2011; Mcintosh et al., 1996; McIntosh & Lobaugh, 2004). In this case, our goal was to identify patterns of physiological measures whose relation with LC contrast differed across age groups. PLS is also well-suited for datasets with highly-correlated predictors (McIntosh et al., 1996), which was relevant since many of our physiological measures were correlated.

Because PLS required the data to be restricted to complete cases, we removed breathing rates from this set of analyses to boost the number of available complete cases to reflect 53 younger and 23 older participants. Then, all measures were centered and normalized. Physiological arousal measures reflecting rest, stress reactivity and stress recovery were stored in a matrix *X*, with rows reflecting individual participants and columns containing the various measures. LC contrast values were stored in a single-column matrix *Y*, with rows reflecting individual participants. The cross-correlation map *R* = *Y*^*T*^*X* was computed for each age group and after arranging the maps in a matrix, the matrix was subjected to singular value decomposition: *R* = *USV^T^*. The resulting left singular vectors (*U*) reflect the LC contrast profiles that best characterized the correlation matrix (also termed “LC saliences”), the right singular vectors (*V*) reflect the physiological profiles that best characterized the correlation matrix (also termed “physiological saliences”), and *S* is a matrix of singular values. The original data *X* and *Y* were then projected onto their respective singular vectors, yielding latent variables of *X* (“physiological scores”; *L_X_* = *XV*) and latent variables of *Y* (“LC scores”; *L_Y_* = *YU*) for each participant.

To test the reliability of identified latent variable(s), a permutation test with 10,000 samples was conducted. This entailed randomly shuffling the rows of *X* but not *Y* and using the distribution of singular values from all permutation samples for testing the null hypothesis of no reliable latent variables (Krishnan et al., 2011; McIntosh & Lobaugh, 2004). Latent variable(s) identified as reliable were then tested for stability through bootstrapping (Krishnan et al., 2011). Specifically, for each of 10,000 bootstrap samples, *X* and *Y* were sampled with replacement, and standard errors were calculated based on physiological saliences across all bootstrap samples. Physiological saliences were divided by their standard errors, yielding a bootstrap ratio for each physiological arousal measure, with each ratio reflecting how much the given arousal measure showed a stable association with LC contrast in the latent variable of interest. Bootstrap ratios with absolute values greater than 2 were considered significantly stable (Krishnan et al., 2011).

The procedure described above was performed three times, once with each LC contrast measure (peak, rostral or caudal) comprising *Y*. PLS correlation analyses were performed in MATLAB using the ‘PLScmd‘ toolbox (Mcintosh et al., 1996). All other analyses were performed in R (Version 4.0.4; R Core Team, 2021). Effect sizes for analyses other than PLS were calculated using the R package ‘effectsizè (Version 0.5; Ben-Shachar et al., 2020) and reported as partial *r*.

## Acknowledgments

We are grateful to the individuals who helped with recruitment and data collection for this project: Sumedha Attanti, Kathryn Cassutt, Katherine Chan, Christine Cho, Paul Choi, Vardui Grigoryan, Ivy Hsu, Ringo Huang, Michael Kwan, Juliana Lee, Jungwon Min, Padideh Nasseri, Lauren Thompson, and Yong Zhang. We thank Sumedha Attanti and Juliana Lee who also performed manual delineation of the LC. This work was supported by National Science Foundation grant number DGE-1842487, and by the National Institute on Aging of the National Institutes of Health under award numbers R01AG057184 and T32AG000037. The content is solely the responsibility of the authors and does not necessarily represent the official views of the National Institutes of Health.

## Footnotes

1 To be consistent with the analyses for older participants, we re-ran all analyses using only values of stress reactivity computed with data from the Stroop task for younger participants. There were no differences in the directionality or significance of results. These results are presented in the Supplementary Results (Section 3).

