## Supplementary Material for "Associations between locus coeruleus MRI contrast and physiological responses to acute stress in younger and older adults"

### Supplementary Methods

#### Section 1. Parameters for HRV analysis

For HRV analysis with the Physionet PhysioNet Cardiovascular Signal Toolbox (Version 1.0.2; Goldberger et al., 2000; Vest et al., 2018), we applied all default settings and parameter values except for the following. Because we performed manual quality control of ECG segments and applied our own post-analysis thresholds for ECG signal quality based on the average sqijw metric for each segment, we set the low-quality threshold for signal quality to be 0 and the rejection threshold to be 1. This led all segments to be analyzed and allowed us to subsequently exclude segments and record detailed metrics on which segments were excluded. In addition, to reduce the number of signals not analyzed, we increased the fraction of data allowed to be missing from a window to 0.20 (missing data occurred after the removal of physiologically implausible r-peaks). Finally, during HRV analysis, we set the window length to the length of the segment being analyzed.

**Section 2. Locations of peak LC intensity**

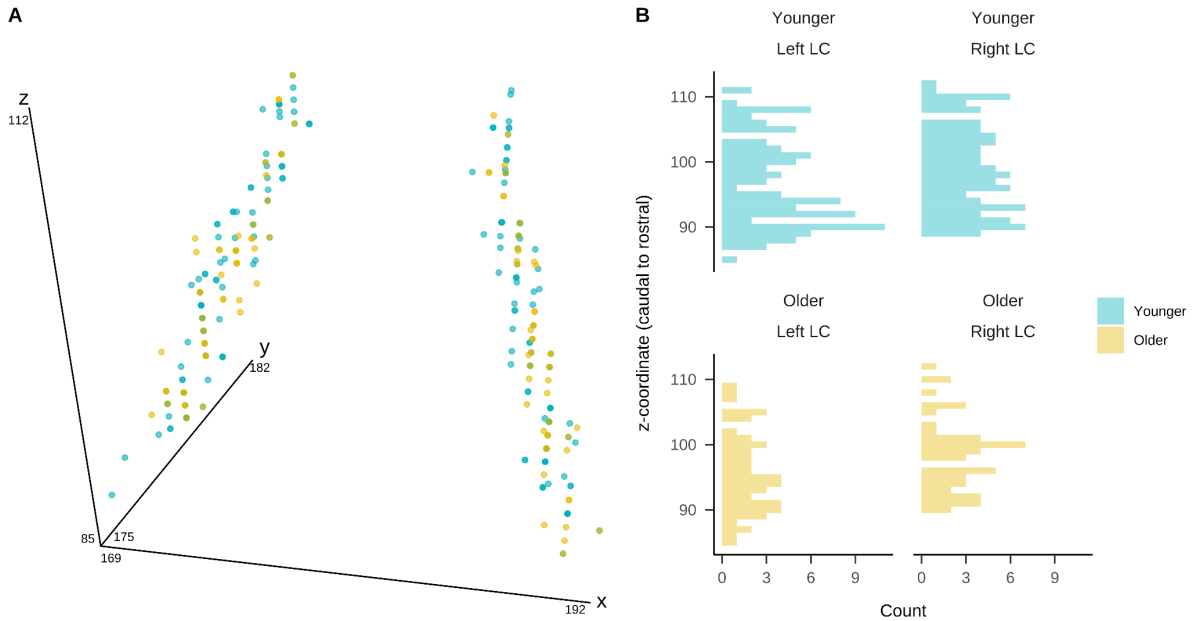


*Figure S1.* (A) 3-dimensional locations of peak LC signal intensity along the LC's rostrocaudal (z-) axis. (B) Histograms depicting distributions of z-coordinates of peak LC signal intensity locations.

**Section 3. Manual LC delineation**

To validate LC intensities determined with the semi-automated approach, we used a manual approach to delineate the LC on native TSE scans. Two trained raters performed the manual delineation protocol described below. On each scan, each rater identified three 2x2 voxel regions of interest (probable LC locations) in each hemisphere in the dorsal pons, near the floor of the fourth ventricle, with maximum signal intensity. This was repeated in each slice in the z-direction where hyperintense voxels near the fourth ventricle were visible. Then, for each hemisphere, the intensity of the maximum-intensity voxel across all slices was extracted. This yielded values of peak left and right LC intensity for each scan, for each rater.

Using intra-class correlation coefficients (two-way models based on absolute agreement), we found high correspondence between LC intensities determined by each rater (left LC: $ICC(3,1) = 0.974$, 95% CI = 0.964 - 0.981, $p < .001$; right LC: $ICC(3,1) = 0.967$, 95% CI = 0.953 - 0.976, $p < .001$). Therefore, we averaged LC intensities for each rater for the purposes of comparing manual intensities with those determined using the semi-automated approach (main text, Section 4.4).

**Section 4. Summary of identified outliers for arousal measures**

**Table S1**

*Summary of outliers identified for average values of each arousal measure.*

| Arousal measure | Age group | N (%) Outliers |
| --- | --- | --- |
| Heart rate | Younger | 3 (1%) |
| Breathing rate | Younger | 2 (0.52%) |
| Systolic blood pressure | Younger | 6 (1.55%) |
| Sympathetic tone (aSKNA) | Younger | 0 (0%) |
| RMSSD | Younger | 3 (1%) |
| log HF power | Younger | 1 (0.33%) |
| log LF power | Younger | 1 (0.33%) |
| Heart rate | Older | 1 (0.82%) |
| Breathing rate | Older | 0 (0%) |
| Systolic blood pressure | Older | 12 (9.52%) |
| Sympathetic tone (aSKNA) | Older | 1 (0.82%) |
| RMSSD | Older | 5 (4.1%) |
| log HF power | Older | 5 (4.1%) |
| log LF power | Older | 5 (4.1%) |

*Note.* Outliers were identified using the mean absolute deviation-median rule, for each age group separately. HF = high-frequency; LF = low-frequency; RMSSD = root mean square of the successive differences.

**Table S2**

*Summary of outliers identified from rolling average values of each arousal measure during the challenge phase.*

| Arousal measure | Age group | N (%) Outliers |
|---|---|---|
| Heart rate | Younger | 275 (1.29%) |
| Breathing rate | Younger | 321 (1.28%) |
| Systolic blood pressure | Younger | 0 (0%) |
| Sympathetic tone (aSKNA) | Younger | 130 (0.6%) |
| RMSSD | Younger | 134 (0.69%) |
| Heart rate | Older | 0 (0%) |
| Breathing rate | Older | 28 (0.62%) |
| Systolic blood pressure | Older | 0 (0%) |
| Sympathetic tone (aSKNA) | Older | 0 (0%) |
| RMSSD | Older | 216 (5.8%) |

*Note.* Before choosing the peak value from 20-second rolling averages of each measure during the challenge phase, we identified outliers using the mean absolute deviation-median rule, for each age group separately. After outlier removal, the peak value during the challenge phase was identified for each measure and participant. For RMSSD, the minimum value across the challenge phase was identified. RMSSD = root mean square of the successive differences.

### Supplementary Results

#### Section 1. Pairwise comparisons of physiological arousal measures by phase, for each age group, during the acute stress task

To test that the acute stress task affected each measure of physiological arousal, we performed planned, pairwise comparisons of each measure for each phase contrast of interest (challenge - baseline, recovery - baseline), for each age group separately. For each set of comparisons, a Bonferroni correction for multiple comparisons was applied. Results of these comparisons for each measure are presented in Tables S3 - S10.

**Table S3**

*Results of pairwise comparisons of estimated marginal means of heart rate during the acute stress induction task, for each phase contrast of interest and age group.*

| Age Group | Contrast | Estimate | SE | *t* | *r* | *p* |
|---|---|---|---|---|---|---|
| Younger | Challenge - Baseline | 4.65 | 0.50 | 9.23 | 0.54 | **<.001** |
| Younger | Recovery - Challenge | -2.91 | 0.51 | -5.71 | -0.37 | **<.001** |
| Younger | Recovery - Baseline | 1.74 | 0.51 | 3.39 | 0.23 | **0.005** |
| Older | Challenge - Baseline | 4.25 | 0.70 | 6.04 | 0.38 | **<.001** |
| Older | Recovery - Challenge | -3.56 | 0.71 | -5.05 | -0.33 | **<.001** |
| Older | Recovery - Baseline | 0.69 | 0.68 | 1.01 | 0.07 | 1.000 |

**Table S4**

*Results of pairwise comparisons of estimated marginal means of breathing rate during the acute stress induction task, for each phase contrast of interest and age group.*

| Age Group | Contrast | Estimate | SE | *t* | *r* | *p* |
|---|---|---|---|---|---|---|
| Younger | Challenge - Baseline | 3.24 | 0.27 | 11.79 | 0.58 | **<.001** |
| Younger | Recovery - Challenge | -3.67 | 0.28 | -13.24 | -0.63 | **<.001** |
| Younger | Recovery - Baseline | -0.43 | 0.27 | -1.56 | -0.09 | 0.717 |
| Older | Challenge - Baseline | 3.84 | 0.42 | 9.08 | 0.48 | **<.001** |
| Older | Recovery - Challenge | -3.45 | 0.42 | -8.15 | -0.44 | **<.001** |
| Older | Recovery - Baseline | 0.39 | 0.42 | 0.94 | 0.06 | 1.000 |

**Table S5**

*Results of pairwise comparisons of estimated marginal means of systolic blood pressure during the acute stress induction task, for each phase contrast of interest and age group.*

| Age Group | Contrast | Estimate | SE | *t* | *r* | *p* |
|---|---|---|---|---|---|---|
| Younger | Challenge - Baseline | 2.59 | 0.84 | 3.07 | 0.19 | **0.014** |
| Younger | Recovery - Challenge | -2.59 | 0.87 | -2.98 | -0.18 | **0.019** |
| Younger | Recovery - Baseline | 0.01 | 0.87 | 0.01 | 0.00 | 1.000 |
| Older | Challenge - Baseline | 4.12 | 1.44 | 2.87 | 0.18 | **0.026** |
| Older | Recovery - Challenge | -7.29 | 1.44 | -5.04 | -0.30 | **<.001** |
| Older | Recovery - Baseline | -3.17 | 1.44 | -2.21 | -0.14 | 0.169 |

**Table S6**

*Results of pairwise comparisons of estimated marginal means of diastolic blood pressure during the acute stress induction task, for each phase contrast of interest and age group.*

| Age Group | Contrast | Estimate | SE | *t* | *r* | *p* |
|---|---|---|---|---|---|---|
| Younger | Challenge - Baseline | 0.86 | 0.56 | 1.53 | 0.10 | 0.769 |
| Younger | Recovery - Challenge | -4.23 | 0.58 | -7.29 | -0.42 | **<.001** |
| Younger | Recovery - Baseline | -3.37 | 0.58 | -5.78 | -0.34 | **<.001** |
| Older | Challenge - Baseline | 1.96 | 0.96 | 2.04 | 0.13 | 0.252 |
| Older | Recovery - Challenge | -4.33 | 0.97 | -4.48 | -0.27 | **<.001** |
| Older | Recovery - Baseline | -2.37 | 0.96 | -2.47 | -0.15 | 0.085 |

**Table S7**

*Results of pairwise comparisons of estimated marginal means of sympathetic tone during the acute stress induction task, for each phase contrast of interest and age group.*

| Age Group | Contrast | Estimate | SE | *t* | *r* | *p* |
|---|---|---|---|---|---|---|
| Younger | Challenge - Baseline | 6.873e-4 | 6.803e-4 | 1.01 | 0.07 | 1.000 |
| Younger | Recovery - Challenge | -1.729e-3 | 6.824e-4 | -2.53 | -0.17 | 0.072 |
| Younger | Recovery - Baseline | -1.042e-3 | 6.991e-4 | -1.49 | -0.10 | 0.825 |
| Older | Challenge - Baseline | 6.861e-4 | 9.596e-4 | 0.72 | 0.05 | 1.000 |
| Older | Recovery - Challenge | -1.923e-3 | 9.687e-4 | -1.99 | -0.13 | 0.290 |
| Older | Recovery - Baseline | -1.237e-3 | 9.330e-4 | -1.33 | -0.09 | 1.000 |

**Table S8**

*Results of pairwise comparisons of estimated marginal means of RMSSD during the acute stress induction task, for each phase contrast of interest and age group.*

| Age Group | Contrast | Estimate | SE | *t* | *r* | *p* |
|---|---|---|---|---|---|---|
| Younger | Challenge - Baseline | -0.07 | 0.04 | -2.09 | -0.14 | 0.228 |
| Younger | Recovery - Challenge | 0.04 | 0.04 | 1.11 | 0.08 | 1.000 |
| Younger | Recovery - Baseline | -0.03 | 0.04 | -0.94 | -0.07 | 1.000 |
| Older | Challenge - Baseline | -0.06 | 0.05 | -1.15 | -0.08 | 1.000 |
| Older | Recovery - Challenge | 0.11 | 0.05 | 2.17 | 0.15 | 0.189 |
| Older | Recovery - Baseline | 0.05 | 0.05 | 1.06 | 0.07 | 1.000 |

**Table S9**

*Results of pairwise comparisons of estimated marginal means of HF power during the acute stress induction task, for each phase contrast of interest and age group.*

| Age Group | Contrast | Estimate | SE | *t* | *r* | *p* |
|---|---|---|---|---|---|---|
| Younger | Challenge - Baseline | -0.12 | 0.08 | -1.52 | -0.10 | 0.781 |
| Younger | Recovery - Challenge | 0.06 | 0.08 | 0.70 | 0.05 | 1.000 |
| Younger | Recovery - Baseline | -0.07 | 0.08 | -0.80 | -0.05 | 1.000 |
| Older | Challenge - Baseline | 0.01 | 0.12 | 0.06 | 0.00 | 1.000 |
| Older | Recovery - Challenge | 0.13 | 0.12 | 1.08 | 0.07 | 1.000 |
| Older | Recovery - Baseline | 0.13 | 0.11 | 1.17 | 0.08 | 1.000 |

**Table S10**

*Results of pairwise comparisons of estimated marginal means of LF power during the acute stress induction task, for each phase contrast of interest and age group.*

| Age Group | Contrast | Estimate | SE | *t* | *r* | *p* |
|---|---|---|---|---|---|---|
| Younger | Challenge - Baseline | -0.21 | 0.08 | -2.70 | -0.18 | **0.044** |
| Younger | Recovery - Challenge | 0.49 | 0.08 | 6.27 | 0.39 | **<.001** |
| Younger | Recovery - Baseline | 0.28 | 0.08 | 3.50 | 0.23 | **0.003** |
| Older | Challenge - Baseline | 0.13 | 0.11 | 1.22 | 0.08 | 1.000 |
| Older | Recovery - Challenge | 0.24 | 0.11 | 2.11 | 0.14 | 0.215 |
| Older | Recovery - Baseline | 0.37 | 0.11 | 3.46 | 0.23 | **0.004** |

**Section 2. Performance on the cognitive challenge tasks**

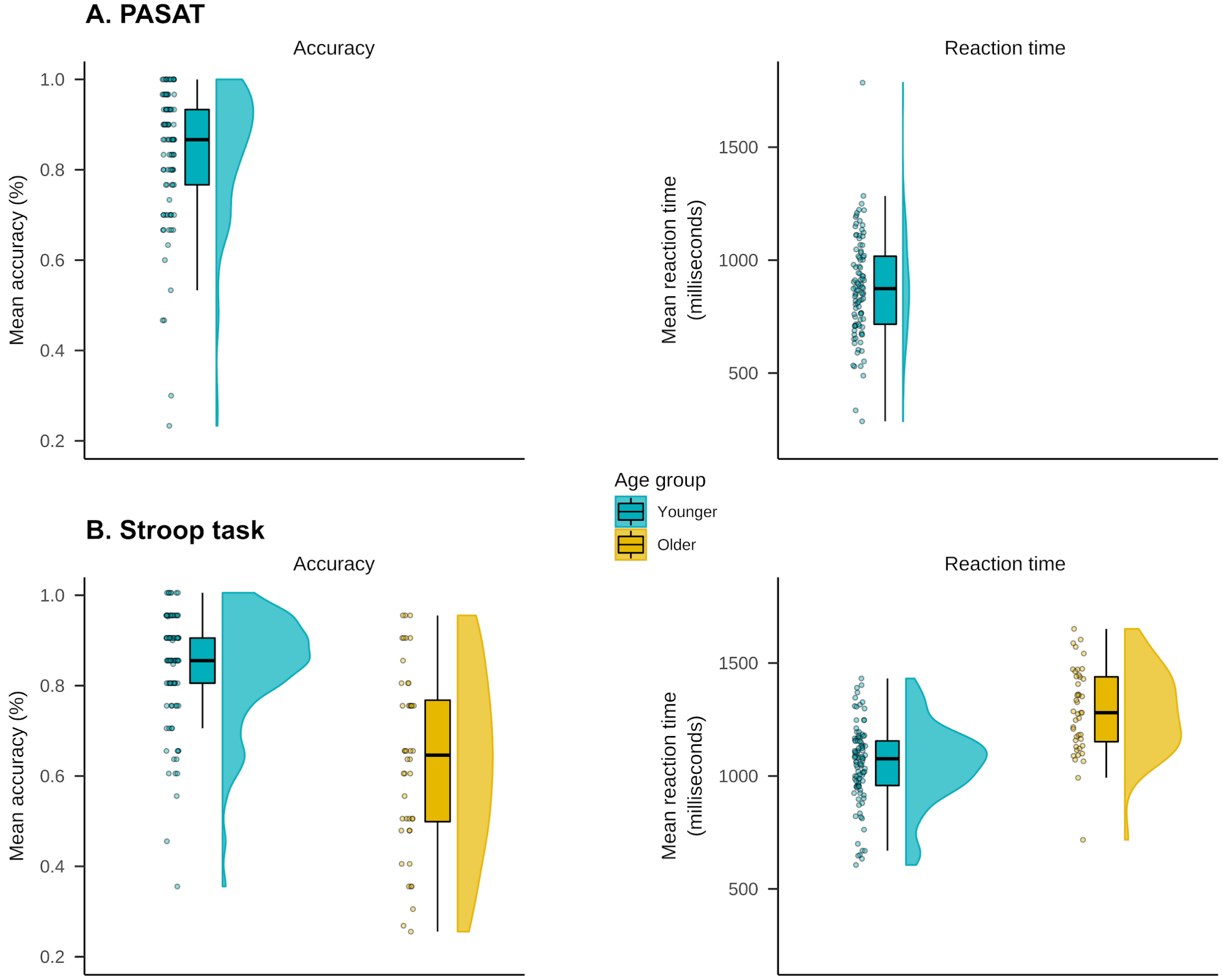


*Figure S2.* Accuracy and reaction times on the Paced Auditory Serial Addition Task (PASAT; A) and Stroop color-word matching task (B).

**Section 3. Results of analyses using only data from the Stroop task for younger participants**

To be consistent with the analyses of older participants, we also performed all LC contrast-arousal analyses using only data from the Stroop task for younger participants. Using this approach, average physiological arousal measures during each phase of the acute stress induction task are shown in Figure S3; in this figure, the challenge phase reflects only the Stroop task for both younger and older participants. After we recomputed values of stress reactivity for younger participants using only data from the Stroop task, we repeated the pairwise Pearson correlation and partial least squares correlation analyses performed in the main text. Pairwise correlation matrices from these analyses are shown in Figure S4. There were no changes to the pattern of significance in the pairwise correlation results for younger participants. In addition, as was the case in the main text, the partial least squares correlation analyses indicated only one marginally reliable latent variable ($p$ = .053) for the analysis of caudal LC contrast and physiological arousal (Figure S5). This latent variable described the same pattern of results as in that in the main text: in older participants, having higher caudal LC contrast was associated with greater systolic blood pressure increases and RMSSD decreases during stress reactivity and higher systolic blood pressure, lower RMSSD, and lower HF power during stress recovery. In summary, using only data from the Stroop task for younger participants did not change the pattern of results.

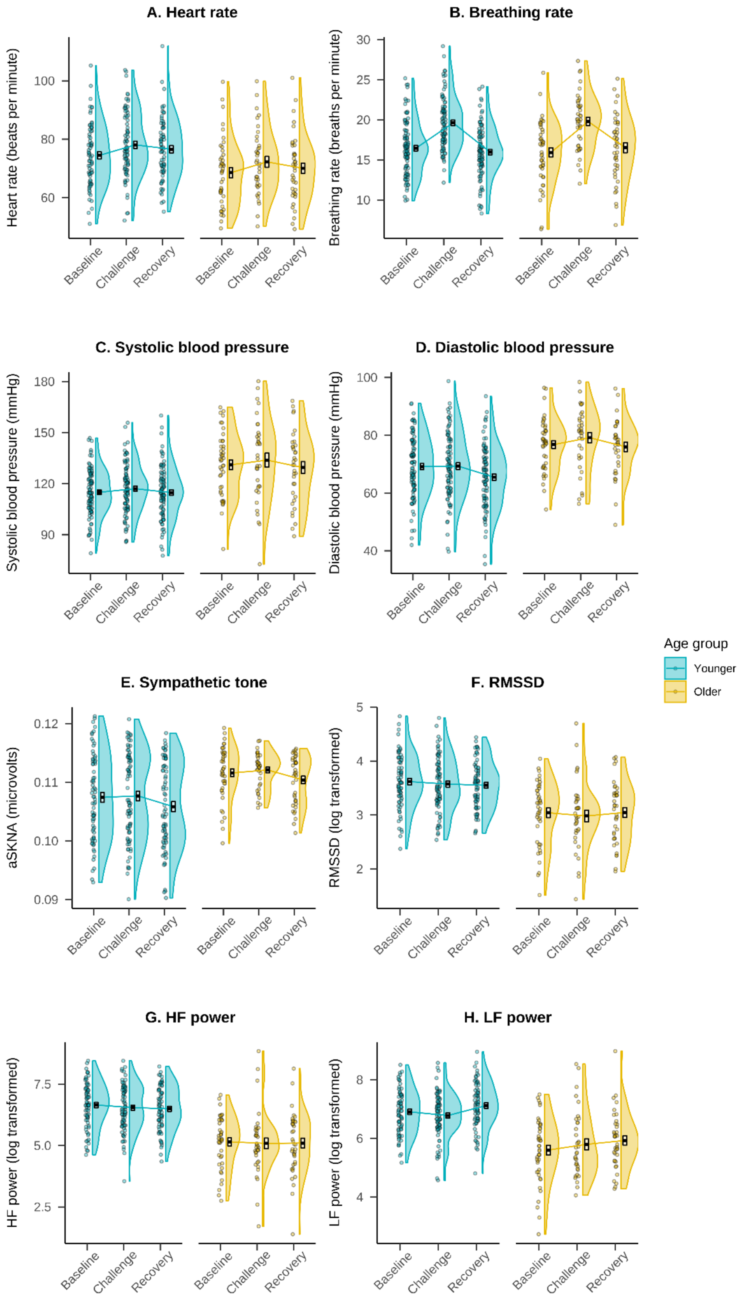


*Figure S3.* Average measures of physiological arousal during each phase of the stress induction protocol, with the challenge phase reflecting only the Stroop task for younger and older participants. Data from older participants is identical to that depicted in Figure 1 in the main text. Crossbars reflect standard errors of the mean. LF = low-frequency; HF = high-frequency; RMSSD = root mean square of the successive differences.

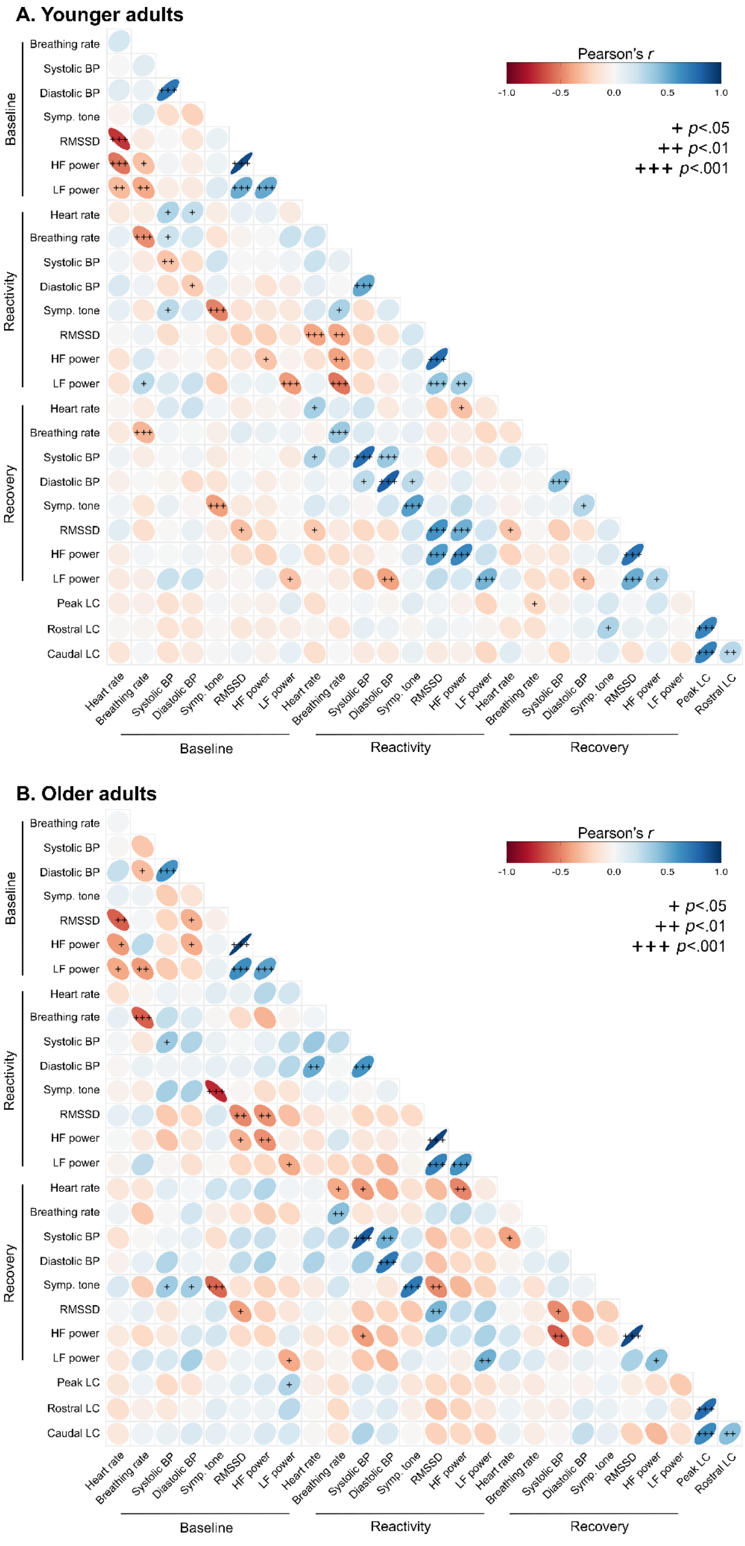


*Figure S4.* Visualization of correlation matrices reflecting pairwise Pearson correlations between LC contrast and arousal during the stress induction protocol. For these analyses, values of stress reactivity were computed using only data from the Stroop task for younger and older participants. Results from older participants are identical to those presented in Figure 4 in the main text. + $p$<.05, ++ $p$<.01, +++ $p$<.001.

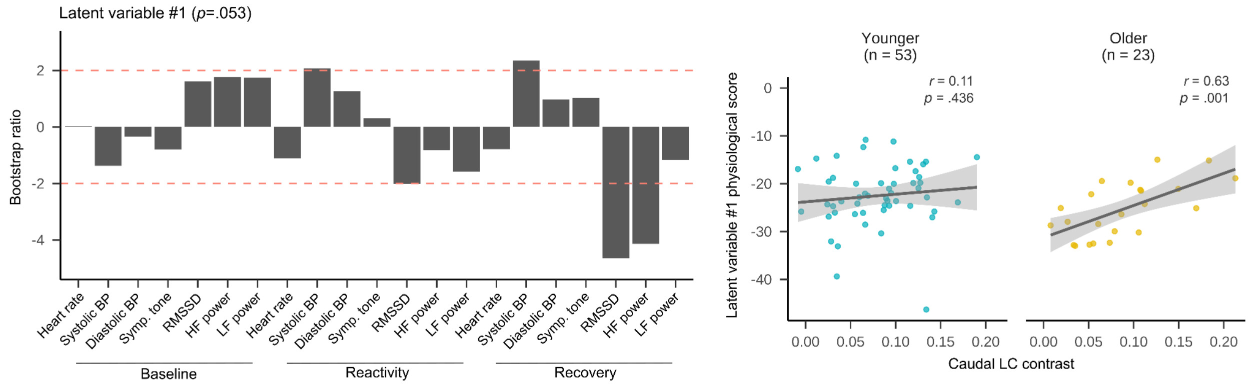


*Figure S5.* Results of partial least squares (PLS) correlation analyses examining the association between caudal LC contrast and physiological arousal during the stress induction task. For these analyses, values of stress reactivity were computed using only data from the Stroop task for younger and older participants. These analyses indicated a marginally reliable latent variable reflecting an association between caudal LC contrast and arousal for older participants. The left panels depict bootstrap ratios which reflect how much each arousal measure contributed to the latent variable (bootstrap ratios with absolute value greater than 2, indicated in red, were considered stable contributors). Right panels depict associations between physiological scores - reflecting the projection of each respective latent variable onto the original physiological arousal data - and LC contrast values.
